# Oxytocin receptor absence reduces selectivity in peer relationships and alters neurochemical release dynamics in prairie voles

**DOI:** 10.1101/2025.05.15.654128

**Authors:** A.B. Black, N. Komatsu, J. Zhao, S.R. Taskey, N.S. Serrano, R. Sharma, D.S. Manoli, M.P. Landry, A.K. Beery

## Abstract

Friendships, or *selective peer relationships*, are a vital component of healthy social functioning in humans, while deficits in these relationships are associated with negative physical and mental health consequences. Like humans, prairie voles are among the few mammalian species that form selective social bonds with both peers and mates, making them an excellent model for the mechanistic investigation of selective social attachment. Here, we explored the role of oxytocin receptors in selective peer attachment using prairie voles lacking a functional oxytocin receptor gene (*Oxtr*^1−/−^). We found that *Oxtr*^1−/−^ animals exhibited significant delays in peer relationship formation compared to wildtype animals. Oxytocin receptor function also contributed to the maintenance of peer bonds, as *Oxtr*^1−/−^ voles displayed reduced relationship stability and lost selective attachments rapidly in a multi-chamber, group-living habitat. *Oxtr*^1−/−^ voles also showed deficits in both general social reward as well as selective social reward for a peer partner over an unfamiliar conspecific. Evoked oxytocin release in the nucleus accumbens was reduced in *Oxtr*^1−/−^ animals compared to their wildtype counterparts, indicating that these voles do not have a compensatory increase in oxytocinergic signaling. Together, these data indicate that oxytocin receptors influence the formation, persistence, and reward value of peer relationships.

## Introduction

Strong and supportive social relationships are crucial for mental well-being, stress resilience, and longevity in social animals^1–5^, while difficulty forming or maintaining relationships is a hallmark of numerous neuropsychiatric disorders^6,7^. Humans form myriad social attachments across the lifespan, including relationships with parents, offspring, romantic partners, group members, and friends. A common feature of these close relationships is *selectivity*, or the preference to spend time with specific social partners over novel individuals. Friendships, referred to here as selective peer relationships, are functionally distinct from reproductive mate relationships, but integral for social health and well-being^5,8–10^. Nevertheless, we lack an understanding of the neural mechanisms that support selective peer relationships.

Across the animal kingdom, the neuropeptide oxytocin has been implicated in various social processes from social recognition and reward to mate bond formation^11–14^. The repeated convergence of complex social behaviors on evolutionarily conserved neural circuits—such as those involving oxytocin signaling—underscores the value of investigating shared neural pathways across diverse taxa as well as unique mechanisms underlying species-specific specializations in social behavior. Growing evidence suggests that oxytocin signaling and receptor distribution play important roles in mediating social interactions between same-sex peers^15–17^, but the role of oxytocin signaling in mediating and reinforcing the development of *selective* social bonds in peer relationships remains limited^18–20^.

Prairie voles have become an essential model organism for understanding the basis of complex social relationships including social monogamy and selective affiliation^21–23^. Unlike most traditional laboratory rodents^24,25^, but similar to humans, prairie voles form selective social bonds with both mates and peers, displaying a robust preference to interact with familiar “partners” over novel “strangers”^26,27^. Foundational pharmacological studies investigating the role of oxytocin signaling in prairie vole mate relationships show that the central and region-specific blockade of oxytocin receptors (OXTRs) impairs mate bond formation^28–30^, while intracranial infusion of oxytocin accelerates the onset of partner preference^30–32^. Though these studies provided seminal insights into understanding the role of oxytocin signaling in relationship formation, early work examined only *mate* attachments.

The novel development of transgenic prairie voles lacking functional OXTRs has paved the way for longitudinal studies of the oxytocin system in social bond formation and other complex social behaviors. Recent studies of OXTR null mutant (*Oxtr*^1−/−^) voles found that functional OXTRs are not required for partner preference formation between mates after one week of cohabitation^33^, but mate relationship formation in *Oxtr*^1−/−^ animals was impaired after *shorter* cohousing intervals that are normally sufficient for bond formation (6 hours for females and 5 days for males)^34^.

These results reinforce early findings from pharmacological approaches, indicating that oxytocin signaling mediates *early* social interactions, accelerates bond formation, and supports the rejection of strangers in reproductive relationships. The concept that oxytocin activity facilitates efficient mate relationship formation parallels existing theories regarding the role of oxytocin in social salience and the amplification of relevant social information^35–37^. Such studies expand our understanding of the role of OXTRs in mate bond formation throughout extended periods of cohabitation but have not explored the effects of OXTR absence on *peer* social relationships.

In the present study, we used *Oxtr*^1−/−^ prairie voles to examine the role of OXTRs in selective peer social relationships. Across multiple behavioral testing paradigms, we found that OXTRs are necessary for early-stage peer bond formation, lasting peer attachment, and selective social reward–specifically in peer contexts. Furthermore, to assess how receptor manipulation affects oxytocin signaling, we employed recently developed synthetic optical probes that enable real-time imaging of extracellular oxytocin dynamics with high spatial and temporal resolution in acute brain slices. Real-time oxytocin imaging revealed that *Oxtr*^1−/−^ animals have reduced evoked oxytocin release in the nucleus accumbens.

## Results

### Peer partner bond formation is delayed in Oxtr^1−/−^ females

We explored how OXTRs impact peer relationship formation by assessing the timeline of peer bond formation in *Oxtr*^1−/−^ and wildtype (*Oxtr*^+/+^; WT) females using partner preference tests (PPT, Figure 1A). WT and *Oxtr*^1−/−^ focal females were pair-housed with a same-sex partner from weaning and tested in adulthood (PPT A: lifetime partner). Next, each focal vole was paired with a new, unrelated peer partner, and two additional PPTs were carried out 24 hours (PPT B) and 7 days (PPT C) into cohabitation to assess new bond formation. We quantified the duration spent in side-by-side contact (huddling) with the partner and the stranger across the three-hour test to characterize social preference. *Oxtr*^1−/−^ females displayed fewer bouts of aggression toward strangers across all three PPTs compared to WT animals (Figure 1B). *Oxtr^−/−^* females formed peer bonds with long-term partners (Figures 2C & 2E) but exhibited significant deficits in the early stages of relationship formation (Figure 2D). There were no significant differences in time spent alone in the center chamber throughout the three tests, or in physiological characteristics such as internal body temperature and adult body weight (Table S1).

**Figure 1.**
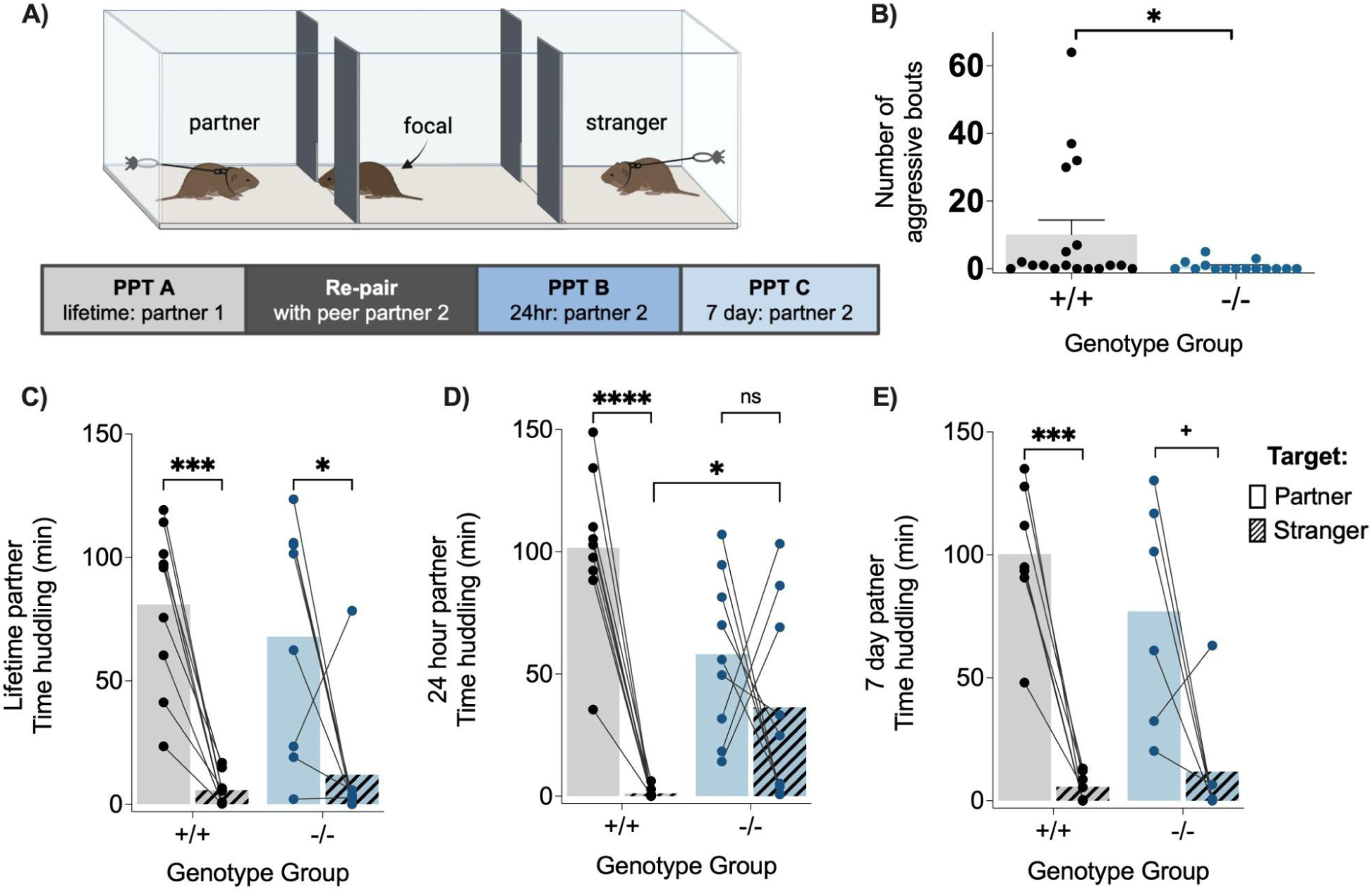
*Oxtr*^1−/−^ females exhibit reduced aggression and delayed peer partner preference formation. **A)** Partner preference test (PPT) apparatus and testing timeline. **B)** Across PPTs, WT (+/+) females were more aggressive than OXTR null mutants (−/−). **C)** Both WT and *Oxtr*^1−/−^ females showed a significant preference to huddle with their lifetime (4+ week) partner versus a stranger. **D)** After 24 hours of cohabitation with the new peer partner, huddling duration was influenced by the interaction of genotype and social target (partner/stranger). WT females showed huddling preference for a 24 hour peer partner while *Oxtr*^1−/−^ voles did not. *Oxtr*^1−/−^ voles also spent more time huddling with the same-sex stranger compared to the WT group. **E)** Both WT and *Oxtr*^1−/−^ females showed a preference for their peer partner after a week of cohabitation. See statistics (model results and pairwise comparisons) reported in Table S1. + = .0527, * = <.05, ** = <.01, *** = <.001, ****<.0001.

**Figure 2.**
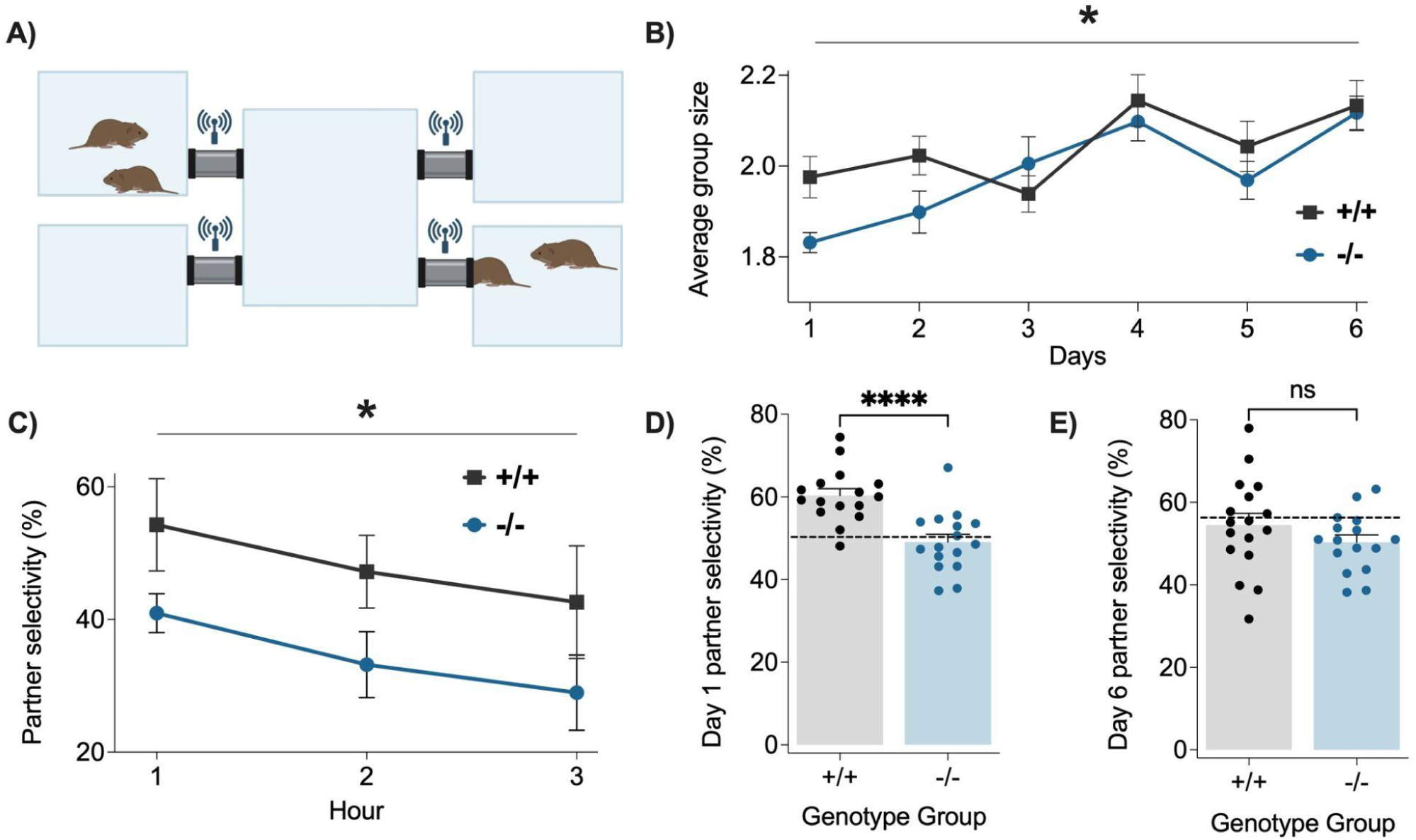
*Oxtr*^1−/−^ females lose track of established relationships and integrate with novel peers in group habitats more quickly than WT females. **A)** 5-chamber habitat with RFID ring antennas at each end of the connecting tubes. **B)** On day 1 and across the week, *Oxtr*^1−/−^ animals cohabited in smaller groups compared to WT animals. **C)** *Oxtr*^1−/−^ voles showed a lower magnitude of partner selectivity (% *partner selectivity = [time with established partner / total time social] * 100*) across the first 3 hours of group living. ‘Genotype’ and ‘test day’ also interacted significantly to affect partner selectivity across the week (see Table S2). **D)** On day one of group cohabitation, *Oxtr*^1−/−^ females spent a lower percentage of their total social time with their partner compared to WT females. The dashed line (51%) indicates the partner selectivity expected by chance on day 1, given cohabitation in the group sizes exhibited on that day (see methods). WT females were significantly more partner selective than expected by chance, whereas *Oxtr*^1−/−^ exhibited no partner selectivity. **E)** By day 6 there was no difference in partner cohabitation by genotype. The dashed line (56%) indicates the partner selectivity expected at random given cohabitation in the group sizes exhibited on that day, and neither group exhibited selective bias toward cohabiting with their original partner. See statistics reported in Table S2.

### Oxtr^1−/−^ females are more gregarious and display less stable peer relationships compared to wildtype females in free-moving groups

We measured the strength and stability of peer bonds within newly formed groups of freely behaving WT and *Oxtr*^1−/−^ females by examining how social patterns changed as pairs became part of larger groups in a semi-naturalistic habitat over 6 days of cohabitation. Each group included two pairs of previously established peer partners (4 voles in total), all of the same genotype. Voles were implanted with RFID tags and placed into a multi-chambered habitat equipped with 8 RFID antennas (Figure 2A) to collect positional data without human disturbance^38^.

*Oxtr*^1−/−^ and WT animals spent the same amount of time in groups, but *Oxtr*^1−/−^ voles formed smaller groups, on average, compared to WT voles (Figure 2B). OXTR function also had a major impact on social selectivity. In the first 3 hours of cohabitation (the duration of a PPT), and across the first day, *Oxtr*^1−/−^ voles spent a smaller proportion of their total social time with their initial partner, suggesting lower partner selectivity compared to WT voles (Figure 2C,D). *Oxtr*^1−/−^ voles were not only less selective than WT voles, they also cohabited with their partner for a duration just below that expected by chance, indicating a complete lack of partner specificity (Figure 2D). By the end of the cohousing period (day 6), there was no difference in partner selectivity between genotype groups, with both groups averaging about half of their total social time with their initial partner (Figure 2E). This convergence in partner selectivity between genotype groups was concurrent with increased time spent in larger groups (i.e. with all voles in the habitat) throughout the week as shown in Figure 2B. Both genotype groups huddled more with new group members after the week of cohousing compared to the pre-test condition, indicating that social relationships developed within each group of voles throughout group living (Figure S1).

### Oxtr^1−/−^ females display deficits in social reward

We quantified selective social motivation in WT and *Oxtr*^1−/−^ females using an operant lever-pressing task coupled with a social choice (Figure 3A). We tested animals in three contexts with various target rewards: 1) peer partner vs stranger, 2) mate partner vs stranger, and 3) mate partner vs non-social reward (novel object). We analyzed the number of rewards of each type (achieved via 4 presses at the respective target lever) across three days of testing as a metric of social motivation.

**Figure 3.**
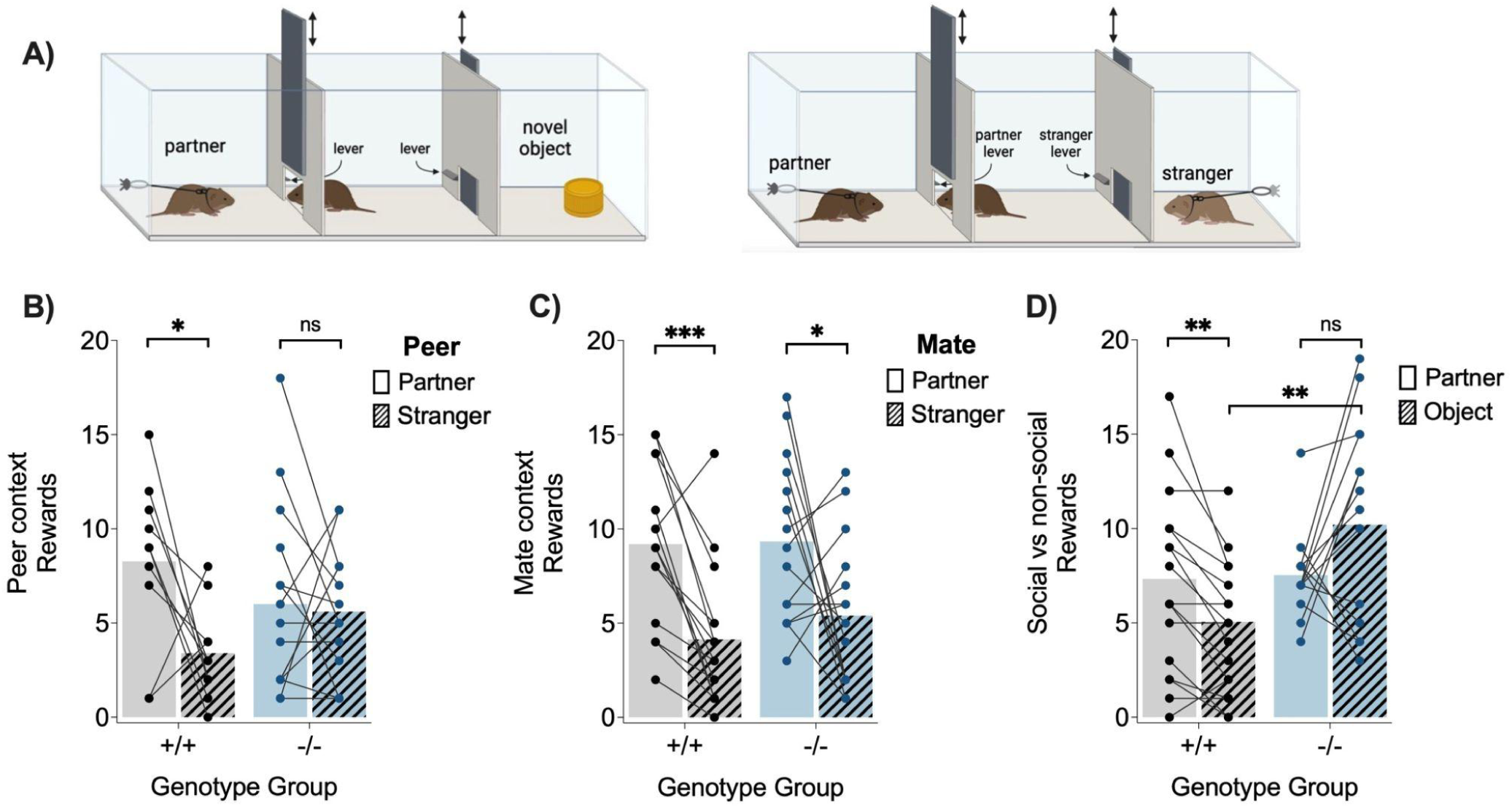
*Oxtr*^1−/−^ voles showed impairments in selective social motivation. **A)** Operant lever pressing choice apparatuses for social and non-social choice contexts. **B)** WT females showed selective motivation for *peer* partners while *Oxtr*^1−/−^ females demonstrated no selective motivation. **C)** Both WT and *Oxtr*^1−/−^ voles displayed selective motivation for their *mate* partner over the stranger. **D)** *Oxtr*^1−/−^ animals showed no selective motivation to access the social (mate partner) over non-social (object) reward, in contrast toWT voles. *Oxtr*^1−/−^ females worked significantly harder to access the novel object compared to WT females. See statistics reported in Table S3.

WT females in peer partnerships were more motivated to access their partner than a stranger, while *Oxtr*^1−/−^ females exhibited no selectivity in peer social reward (Figure 3B). Both WT and *Oxtr*^1-/^ were more motivated to access a mate partner versus a stranger (Figure 3C). However, only WT animals were more motivated to access their mate partner versus a non-social reward. Additionally, *Oxtr*^1−/−^ females showed greater motivation to access this non-social stimulus compared to WT females (Figure 3D). Further behavioral metrics are depicted in Fig S3 (A-C). We found no difference between genotype groups in the rate of reversal learning/extinction of pressing behavior (Fig S3D).

### Oxytocin signaling in the nucleus accumbens is reduced in Oxtr^1−/−^ voles

Given that early bond formation, partner selectivity, and social reward were impaired in *Oxtr*^1−/−^ females, we examined whether oxytocin signaling is enhanced or downregulated in the absence of functional OXTR. We analyzed *ex vivo* oxytocin release kinetics in WT and *Oxtr*^1−/−^ voles following behavioral testing, using recently developed synthetic oxytocin nanosensors ‘NanOx’^39^. NanOx is based on single-walled carbon nanotubes noncovalently functionalized with a single-stranded DNA sequence that selectively modulates fluorescence intensity upon interaction with oxytocin (Figure S4), but not with vasopressin. This synthetic approach provides versatility across species, does not require genetic manipulation^40,41^, and allows for real-time imaging of extracellular oxytocin with high spatiotemporal resolution^39,42^. We imaged electrically evoked oxytocin release in the nucleus accumbens (Figures 4A-C), a region in which OXTR expression and oxytocin signaling have been strongly implicated in pair bonding with mates in prairie voles^28,29,43^.

**Figure 4.**
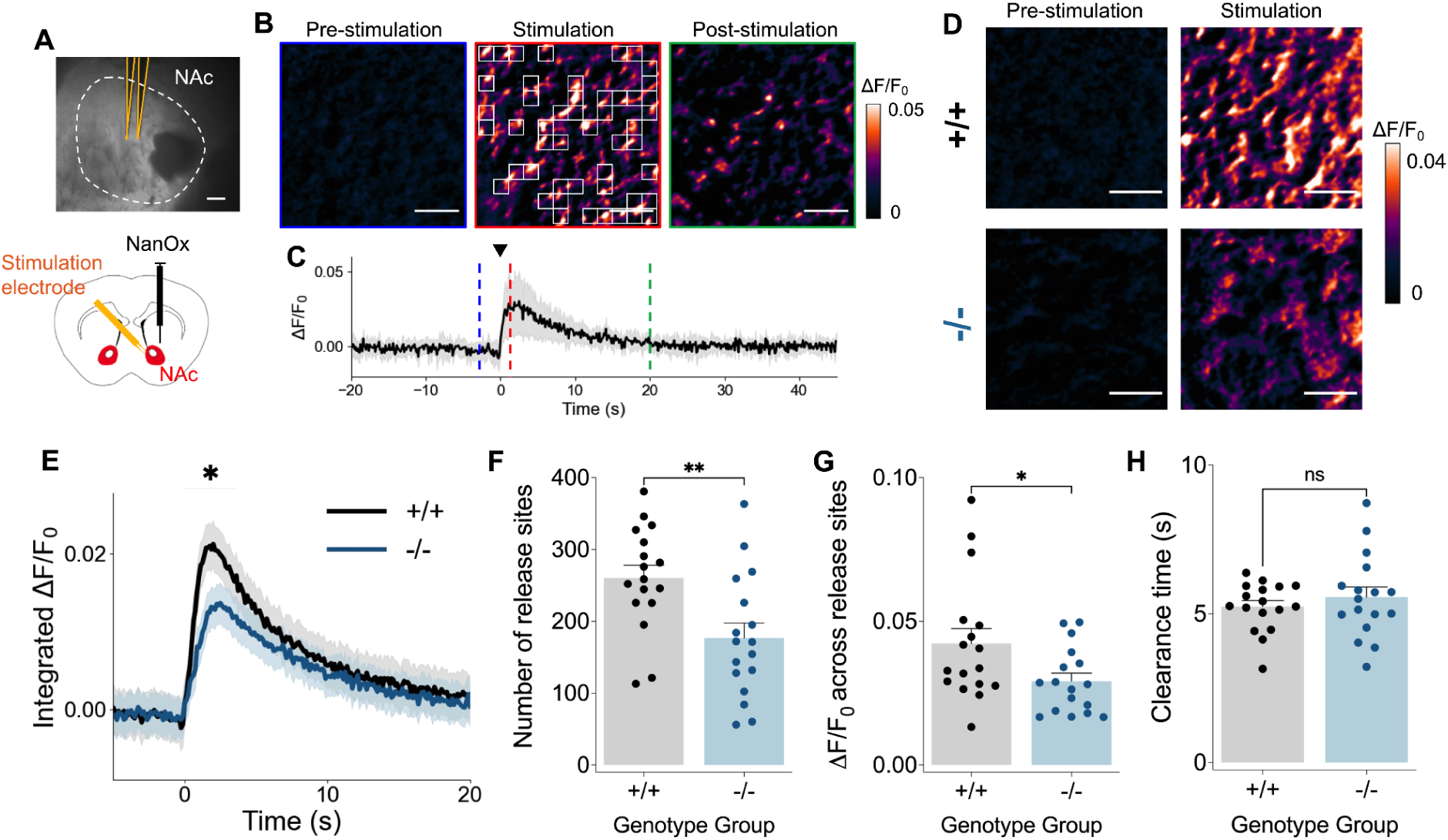
Fluorescence imaging with NanOx reveals reduced oxytocin signaling in the nucleus accumbens of *Oxtr*^1−/−^ voles. **A**) (top) Bright field image and (bottom) schematic showing the nucleus accumbens (NAc) in vole brain slices with stimulation electrodes (yellow). Scale bar represents 200 µm. The extracellular space of brain slices were labeled with NanOx (black). **B**) NanOx ΔF/F_0_ images in nucleus accumbens following 0.5 mA electrical stimulation in standard aCSF with 2 µM quinpirole. Three frames are shown: (left) “pre-stimulation” is the baseline ΔF/F_0_ before electrical stimulation, (center) “stimulation” is immediately after electrical stimulation, where a 6.8 µm by 6.8 µm grid mask was applied and grids with high ΔF/F_0_ were identified as oxytocin release sites (white squares), and (right) “post-stimulation” is after ΔF/F_0_ has returned to baseline. Scale bars represent 20 µm. **C**) Time-course ΔF/F_0_ averaged over release sites in a single field of view (175 µm by 140 µm) in the nucleus accumbens. The solid line is the averaged value of ΔF/F_0_ among identified release sites, and the shadow represents standard deviation (SD). Red, blue, and green dashed lines represent where pre-stimulation, stimulation, and post-stimulation images in (B) are taken from, respectively. **D**) ΔF/F_0_ images of nucleus accumbens (left) before and (right) upon stimulation (left) for (top) WT and (bottom) *Oxtr*^1−/−^ voles. Scale bars represent 20 µm. **E**) Time-course of integrated ΔF/F_0_ in WT and *Oxtr*^1−/−^ voles. The solid line is the averaged value and the shadow is SEM. **F**) The number of release sites identified in the field of view. **G**) ΔF/F_0_ over identified release sites. **H**) Oxytocin clearance time over identified release sites. See statistics reported in Table S4.

We analyzed evoked oxytocin release and quantified several features of oxytocin signaling dynamics (Figures S5 and S6): First, we quantified the number of oxytocin release sites by calculating the regions of interest within the field of view that showed statistically significant increases in NanOx fluorescence upon stimulation. We also quantified oxytocin release by calculating the NanOx fluorescence change (ΔF/F_0_) of the full imaging field of view (integrated ΔF/F_0_), and also of each release site (ΔF/F_0_ across active release sites). Lastly, we quantified the time needed for oxytocin clearance from each release site. We found that *Oxtr*^1−/−^ voles had reduced oxytocin release in the overall imaging field of view (integrated ΔF/F_0_) relative to their wildtype counterparts (Figure 4E). On a subcellular level, *Oxtr*^1−/−^ voles showed fewer oxytocin release sites (Figure 4F), with each release site also exhibiting diminished oxytocin release relative to oxytocin released from WT release sites (Figure 4G). These data suggest both a reduction in the number of oxytocin release sites and amount of oxytocin release across sites in the absence of OXTR. We found no statistically significant difference in the oxytocin clearance rates across genotype (Figure 4H). Motivated by recent studies revealing oxytocin-induced modulation of dopamine pathways in mice and rats^44–47^, we also assessed whether *Oxtr*^1−/−^ mutation impacted accumbal dopamine release in prairie voles using a near-infrared catecholamine nanosensor (nIRCat)^42^. We found that dopamine signaling in *Oxtr*^1−/−^ voles exhibited significantly slower clearance and a trend towards reduced release site density relative to their WT counterparts (Figure S8).

## Discussion

We find that the functional loss of OXTR disrupts selective peer attachment, impacting the formation, stability, and reward value of peer social relationships.

### OXTRs modulate the speed of peer bond formation

Both wildtype and *Oxtr*^1−/−^ voles showed peer partner preference after extended (week-long and lifetime) cohabitation with a peer partner. In contrast, *Oxtr*^1−/−^ females did not form partner preferences after 24 hours of cohabitation—a duration that is sufficient to induce robust partner preferences in wildtype animals^26,32,48^. Abolishing OXTR signaling in female prairie voles thus delayed the formation of selective social relationships with peers, while long-term social bonding remained unaffected. These results reinforce prior pharmacological findings by indicating that oxytocin signaling mediates *early* social interactions and *timely* bond formation^28–30,49–51^. Delays in peer relationship formation parallel deficits found in mate bond formation in *Oxtr*^1−/−^ animals^34^. Together, these data provide further evidence that oxytocin signaling plays a role in the formation of new selective relationships, and acts to amplify relevant social information to efficiently encode salient social contexts^36,37^. While OXTRs play a crucial role in efficient bond formation, the eventual development of a selective attachment must be facilitated by other mechanisms.

### OXTRs are necessary for selectivity in peer groups

Beyond examining the role of OXTRs in dyadic interactions, we probed their role in selective peer social behavior within groups. Unlike monogamous pair bonds with mates, peer relationships in prairie voles are not exclusive and are formed between multiple members of a group^52^. In contrast to wildtype animals, we found that *Oxtr*^1−/−^ voles immediately ‘lost track’ of their original partner in the first 24 hours of free interaction. Indeed, even on the first day of cohousing, *Oxtr*^1−/−^ voles spent no more time with their initial partner than would be expected by chance, indicating a lack of selectivity and an increase in gregarious social behavior. These results suggest that even when *Oxtr*^1−/−^ voles display peer partner preference, the selectivity of their relationships is less stable in broader and more ecologically relevant social settings. It is possible that eventual social attachment seen in partner preference tests is supported by environment and social experience. For instance, wildtype peer partners could facilitate relationship formation through social learning during cohabitation or testing. By using same-genotype cohorts of animals, we parsed out the effects of the *Oxtr*^1−/−^ mutation from social learning or compensation by a wildtype partner, and the loss of selectivity became more obvious in this context. While increased gregariousness may offer advantages such as broader social tolerance and reduced aggression, it may come at the cost of stable relationships and potentially reduce fitness in environments where reliable social bonds support cooperative breeding or resource defense.

Furthermore, OXTRs are known to play an important role in both prosocial and antisocial behaviors in a region- and context-dependent manner^20,53–56^. Deficits in social selectivity found in this assay may arise because the focal animal finds strangers more interesting (reduction in stranger avoidance), or because the animal finds its bonded partner less rewarding (reduced selective social reward) compared to wildtype individuals. These alternatives were assessed by interrogation of the role of OXTRs in selective social reward.

### OXTRs are necessary for selective peer social reward

Social reward plays an important role in reinforcing social relationships^45,57,58^. Importantly, we found no effects of genotype on the voles’ ability to learn, perform, and extinguish the lever-pressing behavior. We found that *Oxtr*^1−/−^ voles displayed impaired selective social motivation to access long-term peer partners, but not mate partners. Moreover, while wildtype animals exhibited stronger motivation for social over non-social reward, *Oxtr*^1−/−^ voles were motivated equally by both reward types and worked harder to access the non-social reward than wildtype animals. These results suggest that OXTRs are not necessary for general reward learning and motivation, but serve a more specific role in social reward and selective motivation for peer partners. Interestingly, intact reward selectivity in *Oxtr*^1−/−^ voles in mate relationships may indicate that mate bonds are more robust and rewarding than peer bonds. Unlike peer relationships, mate relationships are an essential feature of prairie vole mating systems that contribute to reproductive success^59^ (Ophir et al. 2008).

Overall, the loss of social selectivity in *Oxtr*^1−/−^ prairie voles was a deficit observed across diverse ethological contexts, including dyadic and group interactions, various periods of relationship development, and multiple reward contexts, highlighting a critical role for OXTRs in effectively encoding peer relationships. Importantly, although there are no indications of impairments in parental care by OXTR null parents^33^, a constitutive loss of function leads to abnormal neuro-development. Future directions should explore how temporally and spatially precise manipulations of OXTRs in adulthood impact bond formation and selective attachment in peer relationships in a circuit-specific manner.

### Loss of OXTR reduces levels of oxytocin release in the nucleus accumbens

While loss of OXTR does not result in compensatory increase in V1aR abundance^33^, it was unknown whether oxytocin release dynamics were altered in the absence of OXTR function. One possibility was that oxytocin synthesis is upregulated in compensation and signals through vasopressin V1a receptors to elicit similar downstream effects on social behavior. Nonapeptides oxytocin and vasopressin differ by only two of nine amino acids, allowing for substantial affinity at each other’s receptors^60^. Recent studies have noted instances in which these peptides act principally through these alternate pathways^16,19,60–66^. Alternatively, oxytocin signaling could be downregulated in *Oxtr*^1−/−^ voles due to the absence of functional receptors. Thus, we utilized NanOx nanosensors to understand how the *Oxtr*^1−/−^ mutation may alter baseline physiological levels of oxytocin release with very high spatiotemporal resolution.

Evoked oxytocin release was significantly reduced in the nucleus accumbens in *Oxtr*^1−/−^ voles compared to wildtype animals, indicating that oxytocin release is downregulated in the absence of OXTRs. Thus, relationship formation in the absence of OXTR is not accomplished by increased oxytocin acting at V1aR, at least in the nucleus accumbens. The reduction in oxytocin release in the absence of OXTR resulted both from a reduction in number of oxytocin release sites and in the amount of oxytocin released from each site. The reduction in the number of oxytocin release sites may result from fewer oxytocinergic neurons or fewer oxytocinergic projections into the nucleus accumbens due to abnormal neural development. A recent study describing differences in the number of OXT-positive neurons in the PVN of *Oxtr*^1−/−^ voles^34^ supports some of these hypotheses, but further investigation is needed.

Importantly, the coordinated actions of oxytocin and dopamine signaling in the nucleus accumbens reinforce various facets of social behavior across taxa^67^. For instance, both oxytocin and dopamine are known to be major players in social behavior and mate pair-bonding in prairie voles^29,68–71^. Recent studies in meadow voles have also demonstrated an association between greater evoked dopamine release and a social phenotype with greater OXTR density^72,73^. The formation of selective peer social relationships in prairie voles appears not to be dependent on dopamine, although dopamine agonists can facilitate peer bonding^52^, and isolation from either a peer or a mate partner increases D1 receptor binding^74,75^. Here, we found some indications that dopamine signaling in the nucleus accumbens is altered in *Oxtr*^1−/−^ prairie voles. Longer clearance time and a trend toward decreased number of release sites in mutant voles suggest potential changes in dopamine release and reuptake kinetics in the absence of OXTR that may be related to the phenotypic differences in social reward. Future studies are necessary to illuminate potentialmechanisms of OXTR-modulation of dopamine dynamics in peer and mate relationship contexts.

### Conclusions

The present results underscore the role of OXTRs not only in facilitating the timely formation of selective peer relationships but also in maintaining the stability and selectivity of these bonds in broader social contexts. Alongside increased gregariousness observed in groups—characterized by rapid integration with novel conspecifics and loss of peer partner selectivity— *Oxtr*^1−/−^ voles also showed impairments in social reward which may underlie their difficulty maintaining stable, preferential peer relationships. This reduction in social reward, coupled with lower evoked oxytocin release in the nucleus accumbens, suggests that OXTR function is critical not just for forming attachments, but for sustaining the motivational salience of specific peer partners over time. While increased gregariousness and indiscriminate social behavior may promote group cohesion or social flexibility in some contexts, these reductions in social selectivity may compromise the persistence of social bonds and have complex consequences for group dynamics and individual fitness in a socially monogamous and selective species. Together, these findings highlight how OXTR-dependent mechanisms shape the architecture of peer relationships and offer insight into understanding the neural mechanisms that support selective sociality in humans, in whom close social relationships are foundational to mental health, well-being, and resilience.

## Acknowledgements

Jaquesta Adams – NanOx sensor development

Antonella Alaco – Assisted with behavioral testing and scoring

Jolyn Hoang – Assisted with prairie vole genotyping

Alysha Batada, Nicole Chiang, Gautam Naik, Stephan Song – Assisted with operant training

Kelley Power – Assisted with RFID protocols for apparatus set up and data acquisition

UC Berkeley OLAC – Routine animal care

## STAR★Methods

### EXPERIMENTAL MODEL AND STUDY PARTICIPANT DETAILS

Wildtype and *Oxtr*^1−/−^ female prairie voles were bred in-house in a long photoperiod (14 hr light:10 hr dark; lights off at 18:00 PST). Voles were weaned and genotyped at postnatal day (PND) 21 and then separated into same-sex pairs in clear plastic cages with sani-chip bedding and an opaque plastic hiding tube by PND 28. In adulthood (PND 50-80), voles were randomly assigned to one of the three social behavior testing paradigms before use in the oxytocin imaging portion of the study (PND 80-100). There were no significant effects of behavioral testing history on oxytocin release in the imaging experiment.

Only female voles were used to study peer relationships because male prairie voles can show lethal levels of aggression when cohoused with novel, same-sex individuals in adulthood^26^. Furthermore, female peer relationships are more ethologically relevant and are commonly observed in wild prairie voles in contexts such as spontaneous alloparental care^56^. Therefore, experiments that include re-pairing or cohabitation with novel, same-sex animals are restricted to the use of female voles due to animal welfare concerns and interest in biological application to natural social behavior. Both male and female subjects were included in the oxytocin and dopamine imaging experiments to assess possible sex differences in neural signaling. All procedures adhered to federal and institutional guidelines and were approved by the Institutional Animal Care and Use Committee at the University of California, Berkeley.

## METHOD DETAILS

### Genotyping

We collected 1 mm tail samples from animals anesthetized with isoflurane. Tissue samples were lysed according to methods published in Berendzene et al 2023. In short, 200 μL of tail lysis buffer and 10 μL of proteinase K were added to each tube and left to incubate at 56 °C overnight (18–24 hours). Following incubation, samples were briefly spun down and then placed in a 95 °C heat block for 20 minutes to inactivate proteinase K. DNA was subsequently isolated using the Qiagen DNeasy Blood & Tissue Kit, and DNA concentration was quantified using a Nanodrop spectrophotometer. Next, we used polymerase chain reaction and restriction enzyme digestion (Qiagen Taq PCR core kit; XcmI) to establish genotypes according to published protocols (Berendzen et al. 2023). Digested DNA was run on a 2% agarose gel containing SYBR Safe dye in 1X TBE buffer at 140 volts for 45 minutes. The resulting bands were visualized using an Azure Biosystems gel imager.

### Behavioral Testing

#### Partner Preference Test (PPT)

Peer PPTs were carried out using a linear 3-chamber apparatus (Figure 1A) according to previously established methods for preference testing^76,77^. Each behavioral test was recorded and analyzed using automated tracking (idTracker) and scored by a blind observer with custom software (Intervole Timer v1.6; https://github.com/BeeryLab/intervole_timer). We tested the accuracy of automated scoring by comparing it to data from manual scoring on a subset of behavioral tests and found no difference between the two methods in chamber time or huddling time metrics ([*Chamber time:* R² = 0.9933, F(1, 7) = 1031, p = <.0001; *Huddling time:* R² = 0.7484, F(1, 6) = 17.84, p = <.0055). Aggressive bouts were manually scored during the first 20 minutes of recorded tests (PPT 1-3).

In early adulthood (PND 50-70), focal females were tested in the first PPT with a lifetime partner where they had the choice between interacting with a novel “stranger” vole and familiar cagemate “partner” during the 180-minute test. Afterward, each focal vole was separated for three days to prevent aggression upon re-pairing, then placed with a new peer partner for 24-hour cohabitation before the second PPT. Finally, after a week of cohabitation with the new peer partner, the focal females were tested in a third PPT to assess changes in bond formation.

#### Radio-frequency Identification (RFID) Tracking

Free-moving group social dynamics were tracked using RFID tags over a week of undisturbed cohabitation. The group-living habitat is an apparatus composed of five interconnected chambers as previously described by our group^38^. Each chamber is connected by clear acrylic tubes equipped with ring antennas on each side to provide positional tracking of each vole as it moves throughout the habitat. For RFID analysis, raw positional data were collected and processed with Voletron 1.0 (https://github.com/BeeryLab/voletron). Data were analyzed in hour increments for the first three hours and 24-hour intervals across the six days of testing. Social preferences were quantified as time spent with the partner in any size of group/all social time and compared between genotype groups. Each group was also compared to the expected value for partner cohabitation at random (‘expected random’), i.e. if no familiarity preference existed. Expected random values depend on the amount of time individual animals spent in groups of different sizes, as well as the probability of being with the partner by chance in each of those group combinations (i.e. one of three possible pairs involve the partner, two of three possible trios involve the partner, and all possible quads involve the partner). Thus, for each individual, the expected random time with the partner = (⅓*time in pairs)+(⅔*time in trios)+(1*time in quad). These values were averaged across individuals on days 1 and 6 to produce the dashed lines on the graphs (Figure 2D,E), or subtracted from individual partner time to produce supplemental figure S2.

Focal voles were anesthetized with isoflurane and implanted with a subcutaneous RFID tag (BioMark) between the shoulder blades. Focal voles were then placed into the RFID-tracked habitat in groups of four females of the same genotype (*Oxtr*^+/+^ or *Oxtr*^1−/−^). Each trial contained two pairs of two voles (four in total). Each pair of voles lived with their respective partner from weaning, a duration more than sufficient to establish peer partner preference^26^. Since RFID tracking only acquires data for animal position (and lacks behavioral details), we conducted and recorded 10-minute social interaction tests with all four animals before and after the week of group housing in the RFID habitat.

#### Social Operant Conditioning

Training and social testing were conducted according to a previously described operant social choice protocol^78^. Each training apparatus (30.5 cm × 24.1 cm × 21.0 cm) was equipped with a lever, a clicker, and a treat dispenser which released a 20 mg food pellet upon lever press or manual reinforcement. Voles were initially trained to lever press via manual reinforcement in response to lever investigation. Once the vole learned to press independently, it was moved to the second phase of food training at fixed ratio 1. After 3 sequential days of fixed ratio 1, the voles were moved to fixed ratio 4 for 5 days. Voles that learned to lever press according to previously established standards^20^ proceeded to social testing.

Social testing took place in a modular testing apparatus with three chambers (15 cm × 20.5 cm × 13 cm). The center chamber was equipped with two motorized doors and two levers, one of each on either side. A clear plastic tube connects both outside chambers to the center chamber. Prior to social testing, each vole was habituated to the social testing apparatus for 3 days during which the voles learned that each lever opened a door to the corresponding side chamber with a social stimulus rather than a food reward. Before each test, the focal animal was placed into the center chamber where it could press either lever to open the corresponding door allowing access to the reward within the respective chamber for one minute. After one minute the door closes the experimenter moves the focal animal back to the center chamber. Each trial lasted 30 minutes and was recorded for behavioral scoring by a trained observer who was blind to the subject’s experimental group.

Social choice testing occurred in four phases: (1) peer partner versus peer stranger, (2) mate partner versus mate stranger, (3) mate partner versus novel object. Each social testing context was repeated for 3 days. The testing order and novel objects remained the same for both treatment groups. All male partners were castrated and implanted with a testosterone capsule 10-15 days prior to pairing with female focal animals^20^.

### Body weight and temperature

Four solo-housed adults (P60-80) of each genotype were weighed and anesthetized using isoflurane. Each vole was subcutaneously injected with ISO FDX-B thermal sensing PIT tags (BioTherm13, Biomark). Body temperature was monitored for one week in 30-second intervals using an RFID antenna base beneath each cage. The data were processed using ClockLab (Actimetrics/Lafayette Instruments) and averaged for comparison across genotype groups.

### Near-Infared Oxytocin Imaging with NirOX nanosensors

#### Synthesis of ssDNA-SWCNT nanosensors

ssDNA-SWCNT nanosensors were prepared as described previously^39^. Briefly, HiPCo SWCNT slurry (NanoIntegris) and ssDNA (Integrated DNA Technologies) were mixed in a 1.5 mL DNA LoBind tube to a final volume of 1.2 mL with a final NaCl concentration of 10 mM. The mass ratio of DNA to SWCNT was 2:1. The DNA sequence was CCCCCCACGGGACCGCAGATCGAGCCCCCC. The mixture was probe-tip sonicated (Cole-Parmer Ultrasonic Processor, 3-mm tip) in ice-cold water for 30 min at 50% amplitude. The resulting suspensions were centrifuged at 21,000 g for 240 min to pellet unsuspended SWCNT. Nanosensor concentration was calculated by measuring absorbance at 632 nm (NanoDrop One, Thermo Scientific) with an extinction coefficient of ε = 0.036 (mg/L)^−1^ cm^−1^. Nanosensors were diluted to 200 mg/L in 10 mM nucleus accumbensl stored at 4°C.

#### Acute slice preparation and nanosensor labeling

Acute brain slices were prepared following the protocols described in Ref. 24,25. Cutting buffer (119 mM NaCl, 26.2 mM NaHCO3, 2.5 mM KCl, 1mM NaH2PO4, 3.5 mM MgCl4, 10 mM glucose) and aCSF (119 mM NaCl, 26.2 mM NaHCO3, 2.5 mM KCl, 1mM NaH2PO4, 1.3 mM MgCl2, 10 mM glucose, 2 mM CaCl2, all purchased from Sigma-Aldrich) were prepared and bubbled with carbogen gas (oxygen/carbon dioxide 95% O2, 5% CO2, Praxair). The animals were deeply anesthetized via intraperitoneal injection of ketamine (120 mg/kg) and xylazine (24 mg/kg) and transcardially perfused with 5 mL of ice-cold cutting buffer. The brain was subsequently extracted, mounted on a vibratome cutting stage (Leica VT 1000) in the carbogen-bubbled cutting buffer, and cut into 300 µm thick coronal slices with microtome razor blades (Personna, double edge super blades; VWR). Slices were incubated at 37°C for 30 min in oxygen-saturated aCSF in a chamber (Scientific Systems Design, Inc., BSK4) and then transferred to room temperature for 30 min. To label the slice with nanosensors, slices were transferred to a separate incubation chamber (Scientific Systems Design, Inc., BSK2) filled with 5 mL of carbogen-bubbled aCSF at room temperature. Nanosensors were applied to the surface of brain slices with a pipette to a final concentration of 2 mg/L and incubated for 15 min. Slices were rinsed for 5 sec with bubbled aCSF through 3 wells of a 24-well plate to remove unlocalized nanosensors, transferred back to the main chamber, and rested for at least 15 min before imaging.

#### nIR Microscope Design

A custom upright epifluorescence microscope (Olympus, Sutter Instruments) described in greater detail previously (Beyene et al. 2019; Adams et al. 2024) was used to image NanOX and NIRCat fluorescence responses for nanosensors embedded in the extracellular space of acute brain slices. Briefly, a 785 nm excitation laser (OptoEngine LLC, MDL-III-785R-300mW) was used to excite nanosensors, and nanosensor fluorescence was collected with a two-dimensional InGaAs array detector (Raptor Photonics, Ninox 640). Microscope operation was controlled by Micro-Manager Open Source Microscopy Software 42.

#### Electrical stimulation evoked neurochemical imaging with nIR microscope

Carbogen-bubbled aCSF was flown through the microscope chamber with a perfusion pump (World Precision Instruments, Peri-Star Pro) and an aspirator pump (Warner Instruments, Dedicated Workstation Vacuum System) at a rate of 2 mL/min. Imaging chamber temperature was kept at 32°C with an inline heater with feedback control (Warner Instruments, TV-324C). Slices labeled with nanosensors were placed in the chamber with a Tissue harp (Warner Instruments), and a bipolar stimulation electrode (MicroProbes for Life Science, PI2ST30.1A5) was positioned to the targeted field of view, which was adjusted with a 4X objective (Olympus XLFluor 4X/340). Next, NIR imaging was accomplished with a 60X objective (Nikon, CFI Apochromat NIR 60X W) and the imaging field of view was positioned at least 80 µm away from the stimulation electrode. Using Micro-Manager, nIR fluorescence images were acquired at frame rates of 8 frames/s (nominal) for 600 frames, where 1 millisecond of 0.5 mA stimulation was applied at after 200 frames of baseline. This image acquisition was repeated three times with the same field of view, with 5 min of waiting time in between. Note that the electrical stimulation of neurons does not provide cell specificity, potentially leading to the release of other neurochemicals near the electrodes, and NanOx shows response to dopamine while exhibiting selectivity towards other molecules^39^. Therefore, to achieve oxytocin selectivity, 2 µM of quinpirole (Fisher Scientific, 10-611-0), a D2 receptor agonist known to suppress dopamine release^42^, was applied to the chamber through aCSF perfusion and slices were incubated for 15 min before imaging (see Figure S8 for the dataset without quinpirole application). Images were acquired in the same field of view before quinpirole application (three replicates) with 5 min between each replicate stimulation.

#### Image processing and data analysis of nanosensor fluorescence response

Imaging movie files were processed using a custom Python code (https://github.com/NicholasOuassil/NanoImgPro) (schematically summarized in Fig. S5). Integrated ΔF/F_0_ (ΔF/F_0_ for the entire field of view, 175 µm by 140 µm) (Fig. 4D, E, S5A and S5D) was calculated as ΔF/F_0_ = (F-F_0_) /F_0_, where F_0_ is the average intensity for the first 5% of frames and F is the dynamic fluorescence intensity from the entire field of view (Fig. S5B) and averaged over three stimulation replicates. Next, a 25×25 pixel (corresponding to 6.8 µm by 6.8 µm) grid mask was applied to the image stack to minimize bias and improve stack processing time, and then a median filter convolution within each grid was calculated. For each grid square, ΔF/F_0_ was calculated, and regions of interest (ROI) were identified if the F at the time of stimulation (200 frames) was 3 standard deviations above the baseline F_0_ activity (Fig. S5E). Subsequently, the number of ROIs identified from three stimulation replicates was averaged and called the number of release sites. Lastly, ΔF/F_0_ was averaged over all identified ROIs, corresponding to the ΔF/F_0_ across active release sites (Fig. S5F). ΔF/F_0_ for the entire field of view can also be obtained by applying a 640×640 pixel to the Python code above. The grid size dependence and the threshold value dependence of each parameter were evaluated to identify optimal values for each parameter (Figures S6A and C).

## QUANTIFICATION AND STATISTICAL ANALYSIS

All data are shown as means±standard error of the mean (SEM). Statistical significance is determined by an α of 0.05. Analysis of Variance (ANOVA) was used for comparing multiple groups and general linear mixed models (GLMM) were utilized for experiments in which subjects underwent repeated days of testing. Significant effects were followed up by post hoc comparisons. All statistical tests, effects, and sample sizes for primary and supplementary figures are articulated in the supplementary data tables or figure captions, respectively. GraphPad Prism 10 or JMP Pro 17 were used for all data analysis.

## KEY RESOURCES TABLE

### TABLE FOR AUTHOR TO COMPLETE

***<u>Please do not add custom subheadings.</u>*** *If you wish to make an entry that does not fall into one of the subheadings below, please contact your handling editor or add it under the “other*” subheading*. **<u>Any subheadings</u> <u>not relevant to your study can be skipped.</u>** (**NOTE:** references should be in numbered style, e.g., Smith et al.*^1^)

#### Key resources table

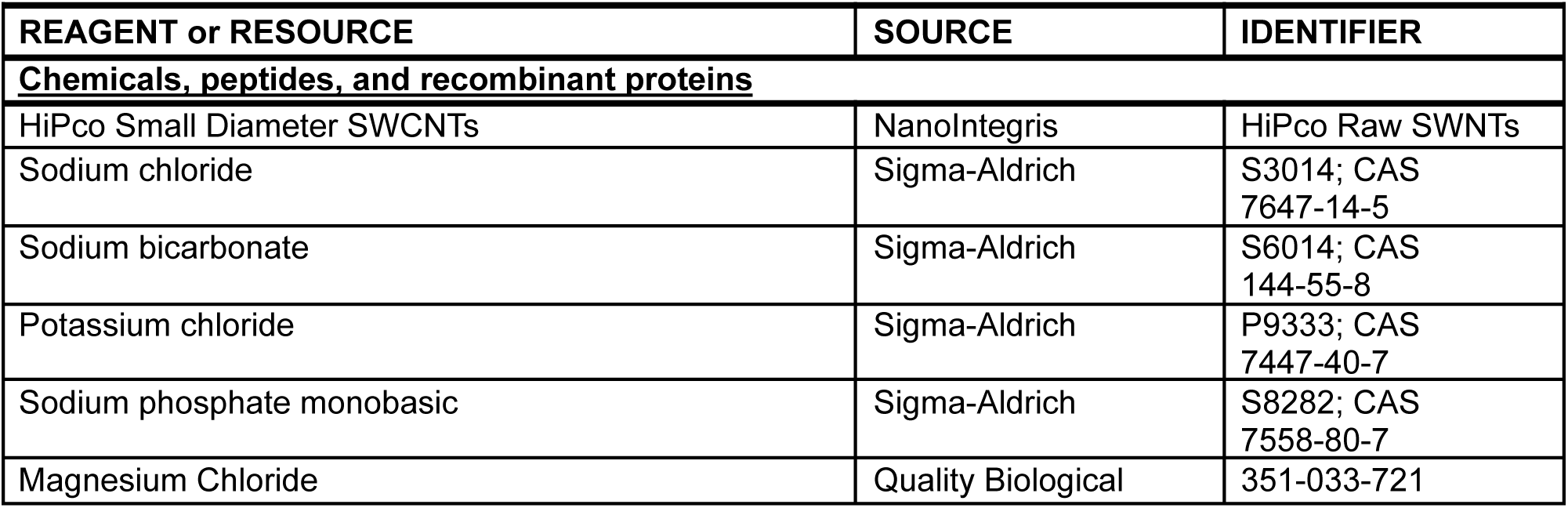
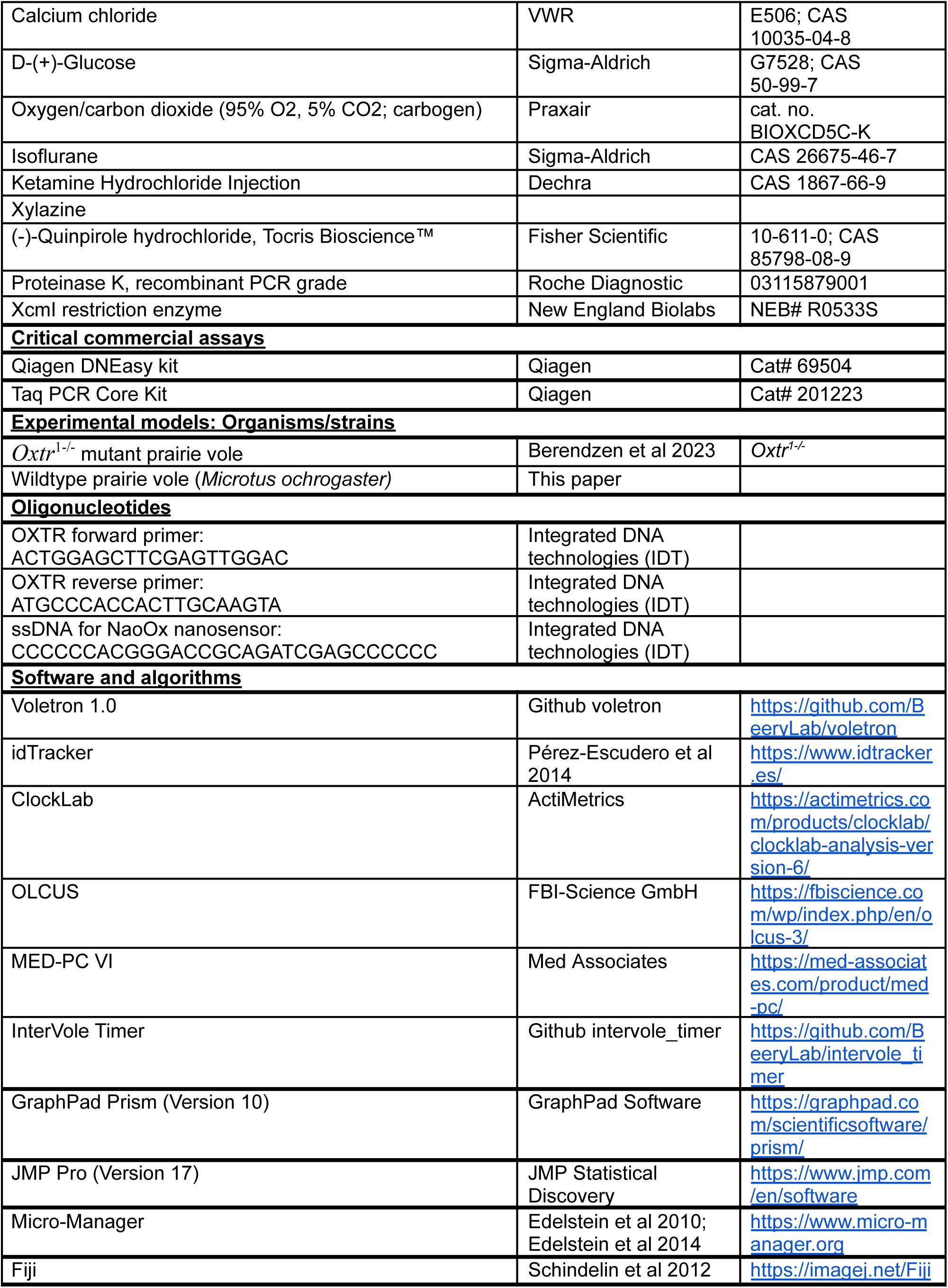
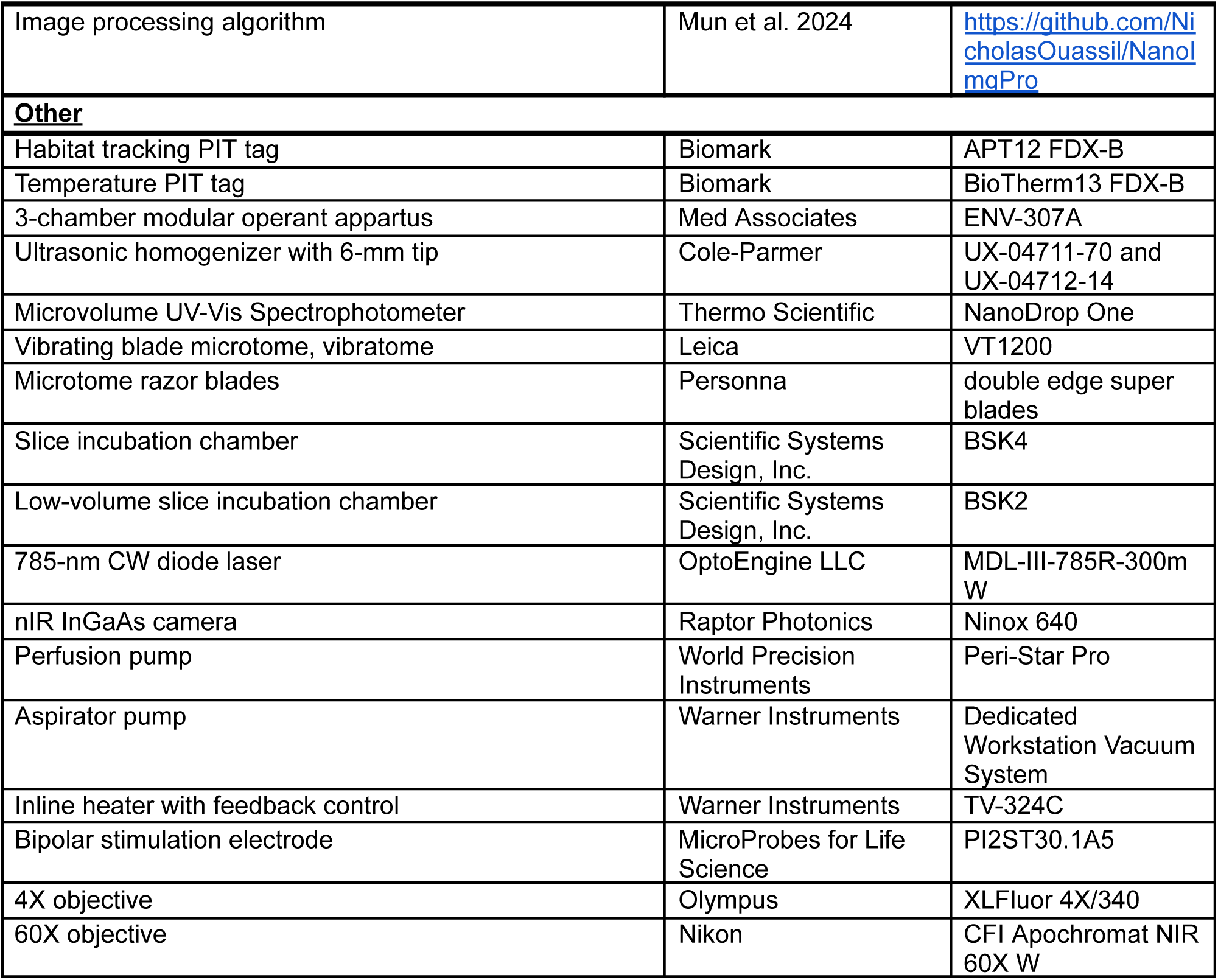

---

**Table S1.** Statistics for Experiment 1.

| Stat Test | Statistic | P value |
| --- | --- | --- |
| **<u>(1B) Sample size</u>** | Animals: 6 +/+; 5 -/-<br>Behavioral tests: 18 +/+; 15 -/- | N/A |
| **1B) GLMM -2LL: 240**<br>Genotype [+/+;-/-]<br>Random: Subject | Variance estimate = -1.0326<br>F(1,14.4) = 5.1921<br>Variance estimate = 2.4439 | 0.0384<br>0.0322 |
| **<u>(1C) Sample size</u>** | Animals: 9 +/+; 8 -/- | N/A |
| **1C) 2x2 ANOVA**<br>Target [partner; stranger]<br>Genotype [+/+;-/-]<br>Target X genotype | F(1,30) = 36.4896<br>F(1, 30) = 0.09641<br>F(1,30) = 0.7962 | <.0001<br>0.7583<br>0.3793 |
| **1C) Paired t test**<br>Partner vs stranger (+/+)<br>Partner vs stranger (-/-) | t(8) = 6.909<br>t(7) = 2.489 | 0.000123<br>0.04166 |
| **<u>(1D) Sample size</u>** | Animals: 10 +/+; 9 -/- | NA |
| **1D) 2x2 ANOVA**<br>Target [partner; stranger]<br>Genotype [+/+;-/-]<br>Target X genotype | F(1,32) = 36.4700<br>F(1,32) = 0.1706<br>F(1,32)=15.1619 | <.0001<br>0.6823<br>0.0005 |
| **1E) Paired t test**<br>Partner vs stranger (+/+)<br>Partner vs stranger (-/-) | t(8) = 9.532<br>t(8) = 0.9275 | 0.000012<br>0.3808 |
| **<u>(1E) Sample size</u>** | 7 +/+; 6 -/- | NA |
| **1E) 2x2 ANOVA**<br>Target [partner; stranger]<br>Genotype [+/+;-/-]<br>Target X genotype | F(1,22) = 48.0815<br>F(1, 22) = 0.5518<br>F(1,22)=0.2162 | <.0001<br>0.4654<br>0.2162 |
| **1E) Paired t test**<br>Partner vs stranger (+/+)<br>Partner vs stranger (-/-) | t(6) = 8.491<br>t(5) = 2.527 | 0.000146<br>0.05270 |
| **<u>(PPT center time) Sample size</u>**<br>**Welch's t test**<br>Center time +/+ vs -/- | Animals: 10 +/+; 9 -/-<br>Behavioral tests: 25 +/+, 23 -/-<br>t(36.37) = 1.321 | NA<br>0.1949 |
| **<u>(Body temp) Sample size</u>**<br>**Student's t test**<br>Body temp +/+ vs -/- | Animals: 4 +/+; 4 -/-<br>t(6) = 1.259 | NA<br>0.2548 |
| **<u>(Body mass) Sample size</u>**<br>**Student's t test**<br>Body mass +/+ vs -/- | Animals: 8 +/+; 9 -/-<br>t(15) = 0.2304 | NA<br>0.8209 |

**Table S2.**
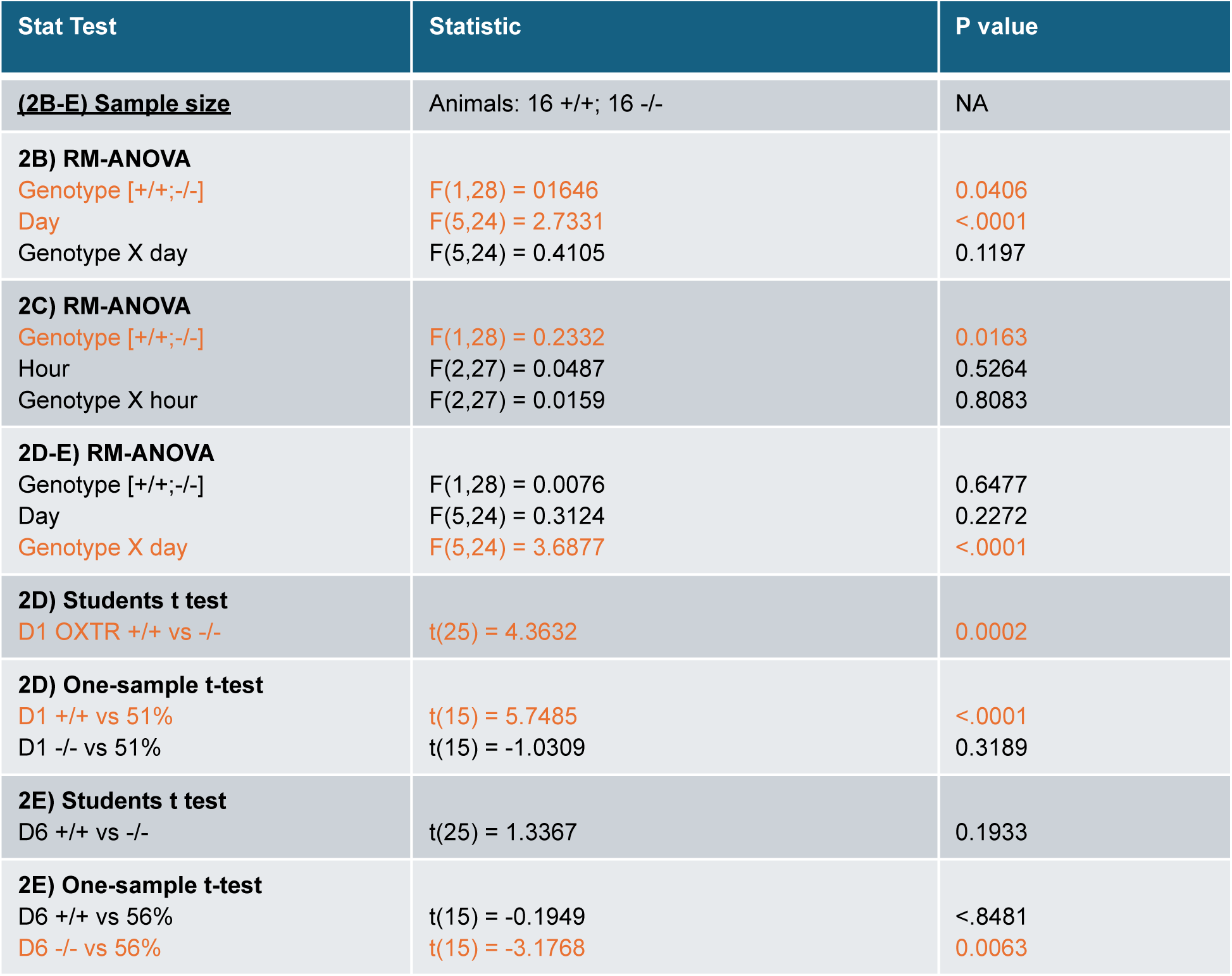
Statistics for Experiment 2.

**Table S3.**
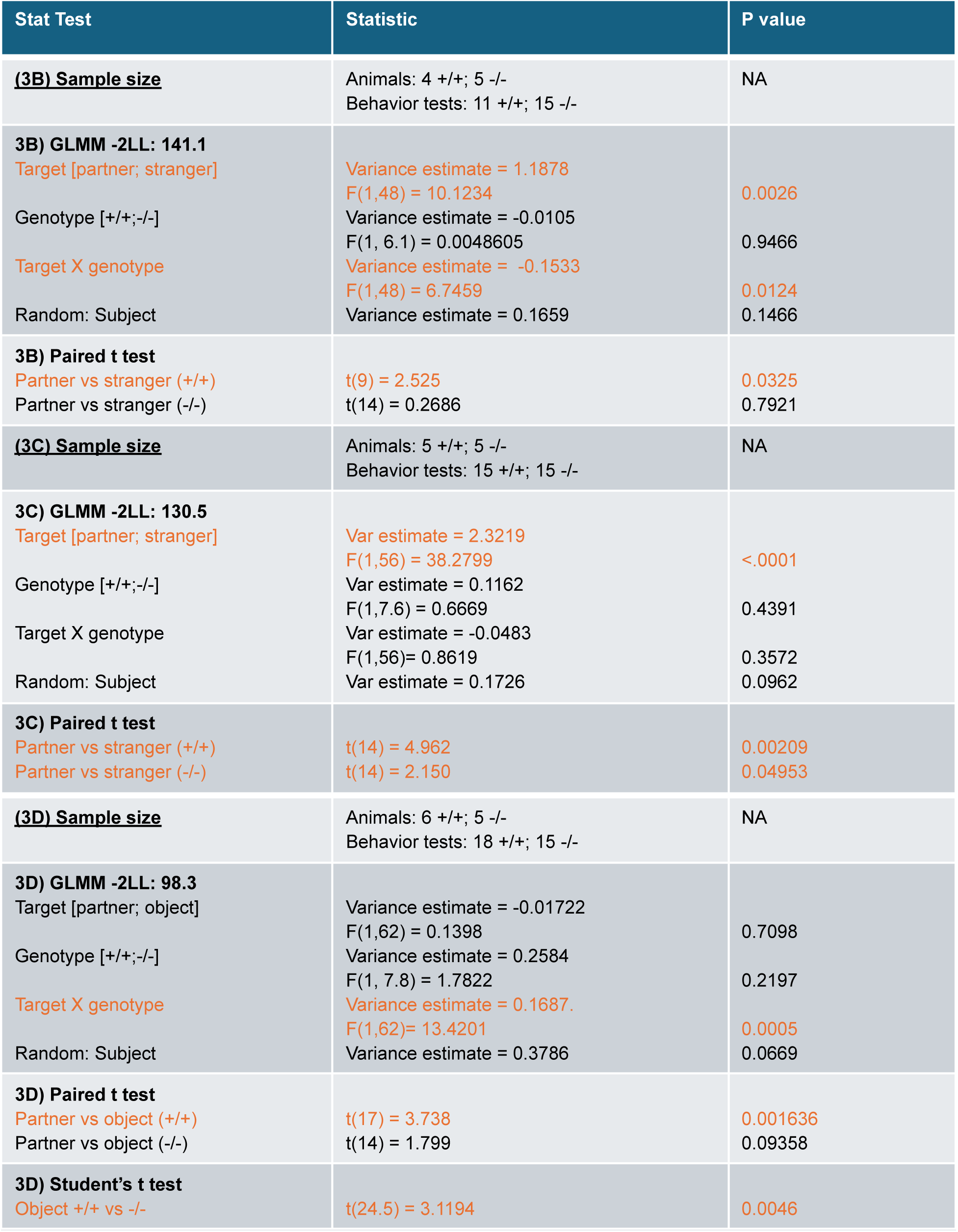
Statistics for Experiment 3.

| Stat Test | Statistic | P value |
| --- | --- | --- |
| **(3B) Sample size** | Animals: 4 +/+; 5 -/-<br>Behavior tests: 11 +/+; 15 -/- | NA |
| **3B) GLMM -2LL: 141.1**<br>Target [partner; stranger] | Variance estimate = 1.1878<br>F(1,48) = 10.1234 | 0.0026 |
| Genotype [+/+;-/-] | Variance estimate = -0.0105<br>F(1, 6.1) = 0.0048605 | 0.9466 |
| Target X genotype | Variance estimate = -0.1533<br>F(1,48) = 6.7459 | 0.0124 |
| Random: Subject | Variance estimate = 0.1659 | 0.1466 |
| **3B) Paired t test**<br>Partner vs stranger (+/+) | t(9) = 2.525 | 0.0325 |
| Partner vs stranger (-/-) | t(14) = 0.2686 | 0.7921 |
| **(3C) Sample size** | Animals: 5 +/+; 5 -/-<br>Behavior tests: 15 +/+; 15 -/- | NA |
| **3C) GLMM -2LL: 130.5**<br>Target [partner; stranger] | Var estimate = 2.3219<br>F(1,56) = 38.2799 | <.0001 |
| Genotype [+/+;-/-] | Var estimate = 0.1162<br>F(1,7.6) = 0.6669 | 0.4391 |
| Target X genotype | Var estimate = -0.0483<br>F(1,56)= 0.8619 | 0.3572 |
| Random: Subject | Var estimate = 0.1726 | 0.0962 |
| **3C) Paired t test**<br>Partner vs stranger (+/+) | t(14) = 4.962 | 0.00209 |
| Partner vs stranger (-/-) | t(14) = 2.150 | 0.04953 |
| **(3D) Sample size** | Animals: 6 +/+; 5 -/-<br>Behavior tests: 18 +/+; 15 -/- | NA |
| **3D) GLMM -2LL: 98.3**<br>Target [partner; object] | Variance estimate = -0.01722<br>F(1,62) = 0.1398 | 0.7098 |
| Genotype [+/+;-/-] | Variance estimate = 0.2584<br>F(1, 7.8) = 1.7822 | 0.2197 |
| Target X genotype | Variance estimate = 0.1687<br>F(1,62)= 13.4201 | 0.0005 |
| Random: Subject | Variance estimate = 0.3786 | 0.0669 |
| **3D) Paired t test**<br>Partner vs object (+/+) | t(17) = 3.738 | 0.001636 |
| Partner vs object (-/-) | t(14) = 1.799 | 0.09358 |
| **3D) Student's t test**<br>Object +/+ vs -/- | t(24.5) = 3.1194 | 0.0046 |

**Table S4.** Statistics for Experiment 4.

| Stat Test | Statistic | P-value |
| --- | --- | --- |
| **(4E-H) Sample size** | Animals: 10 +/+ (6F; 4M); 10 -/- (7F; 3M)<br>Slices: 17 +/+ (9F; 8M); 17 -/- (11F; 6M) | N/A |
| **4E) 2x2 ANOVA**<br>Sex [female; male]<br>Genotype [+/+;-/-]<br>Sex X genotype | F(1, 30) = 0.4871<br>F(1, 30) = 2.983<br>F(1, 30) = 1.127 | 0.4906<br>0.0944<br>0.2969 |
| **4E) Students t test**<br>+/+ vs -/- (both sex) | t(32) = 2.060 | 0.0477 |
| **4F) 2x2 ANOVA**<br>Sex [female; male]<br>Genotype [+/+;-/-]<br>Sex X genotype | F(1, 30) = 0.1393<br>F(1, 30) = 8.5807<br>F(1, 30) = 0.0266 | 0.7116<br>0.0064<br>0.8715 |
| **4F) Students t test**<br>+/+ vs -/- (both sex) | t(32) = 3.074 | 0.0043 |
| **4G) 2x2 ANOVA**<br>Sex [female; male]<br>Genotype [+/+;-/-]<br>Sex X genotype | F(1, 30) = 0.4865<br>F(1, 30) = 4.0054<br>F(1, 30) = 0.0637 | 0.4909<br>0.0545<br>0.8025 |
| **4G) Students t test**<br>+/+ vs -/- (both sex) | t(32) = 2.231 | 0.0328 |
| **4H) 2x2 ANOVA**<br>Sex [female; male]<br>Genotype [+/+;-/-]<br>Sex X genotype | F(1, 30) = 0.0265<br>F(1, 30) = 1.0370<br>F(1, 30) = 0.8767 | 0.8718<br>0.3167<br>0.3566 |
| **4H) Students t test**<br>+/+ vs -/- (both sex) | t(32) = 0.8063 | 0.4260 |

**S1.**
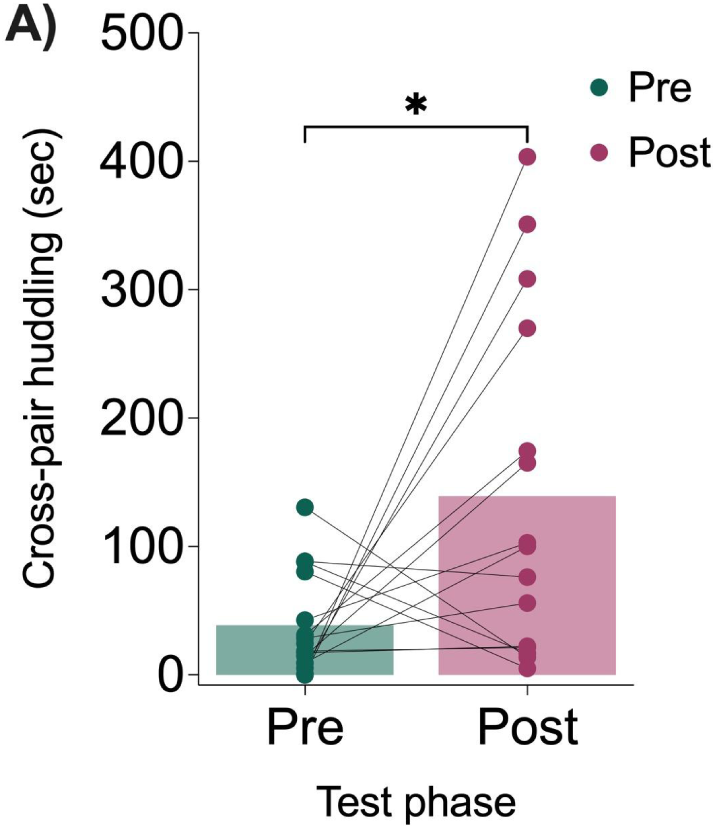
Differences in behavior during pre- and post-group living social interaction. **A)** Individuals spent significantly more time huddling in groups containing one or both unfamiliar ‘cross-pair’ animals (in a pair, trio, or quad) following a week of cohabitation in the group living habitat (pooled across genotype groups). Data (mean ± SEM) from 5 (pre-test) and 5 (post-test) cohorts of 4 voles. Paired t-test t(14) = 2.405, p = 0.0306.

**S2.**
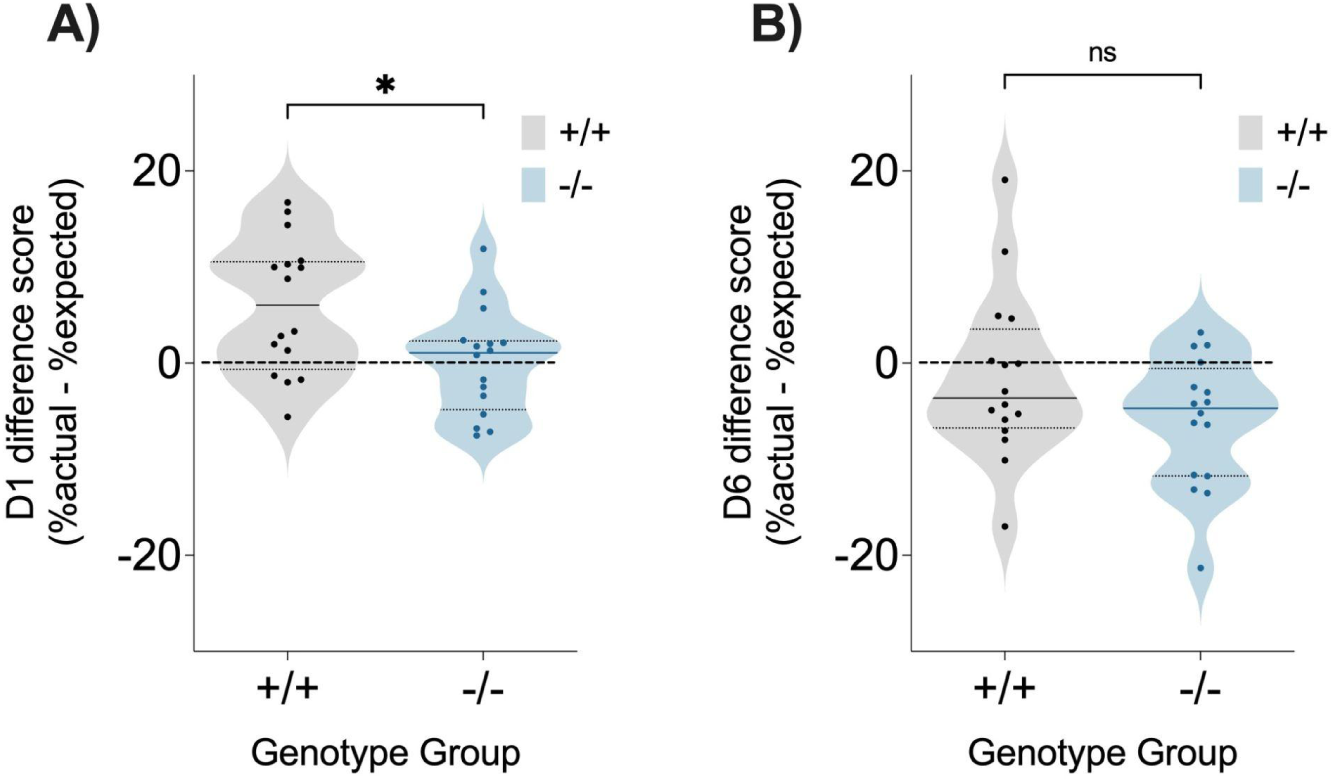
OXTR null mutants lack partner selectivity when novel animals are added to a group context. In the main manuscript (Figure 2) partner time is compared to an expected random value computed using average times across all subjects. Here, difference scores use the expected random value for each individual based on their unique duration of cohabitation in pairs, trios, and quads on that day as: ⅓*(time in pairs)+⅔*(time in trios)+1*(time in quad) (described further in methods). **A)** *Oxtr*^1−/−^ voles a complete lack of partner selectivity across the first 24 hours (one sample t test *Oxtr*^1−/−^ vs 0: t(15) = 0.0487, p = 0.9618). In contrast, the difference scores of WT voles were significantly greater than 0 (one sample t test *Oxtr*^1+/+^ vs 0: t(15) = 3.4256, p = 0.0038), indicating the expected partner selectivity. Student’s t-test between groups t(30) = 2.660, p = 0.0124. **B)** By day 6, there was no difference between genotype groups in partner selectivity (student’s t-test between groups t(30) = 1.618, p = 0.1162). The difference scores for both groups appeared near or below zero, indicating an absence of selectivity for their initial partner after a week of group cohabitation (one sample t test WT vs 0: t(15) = -0.7282, p = 0.4777; *Oxtr*^1−/−^ vs 0: t(15) = -3.5688, p = 0.0028). Data (median ± quartiles) from 16 (WT) and 16 (*Oxtr*^1−/−^) voles, dashed line indicates the expected value at random (0).

**S3.**
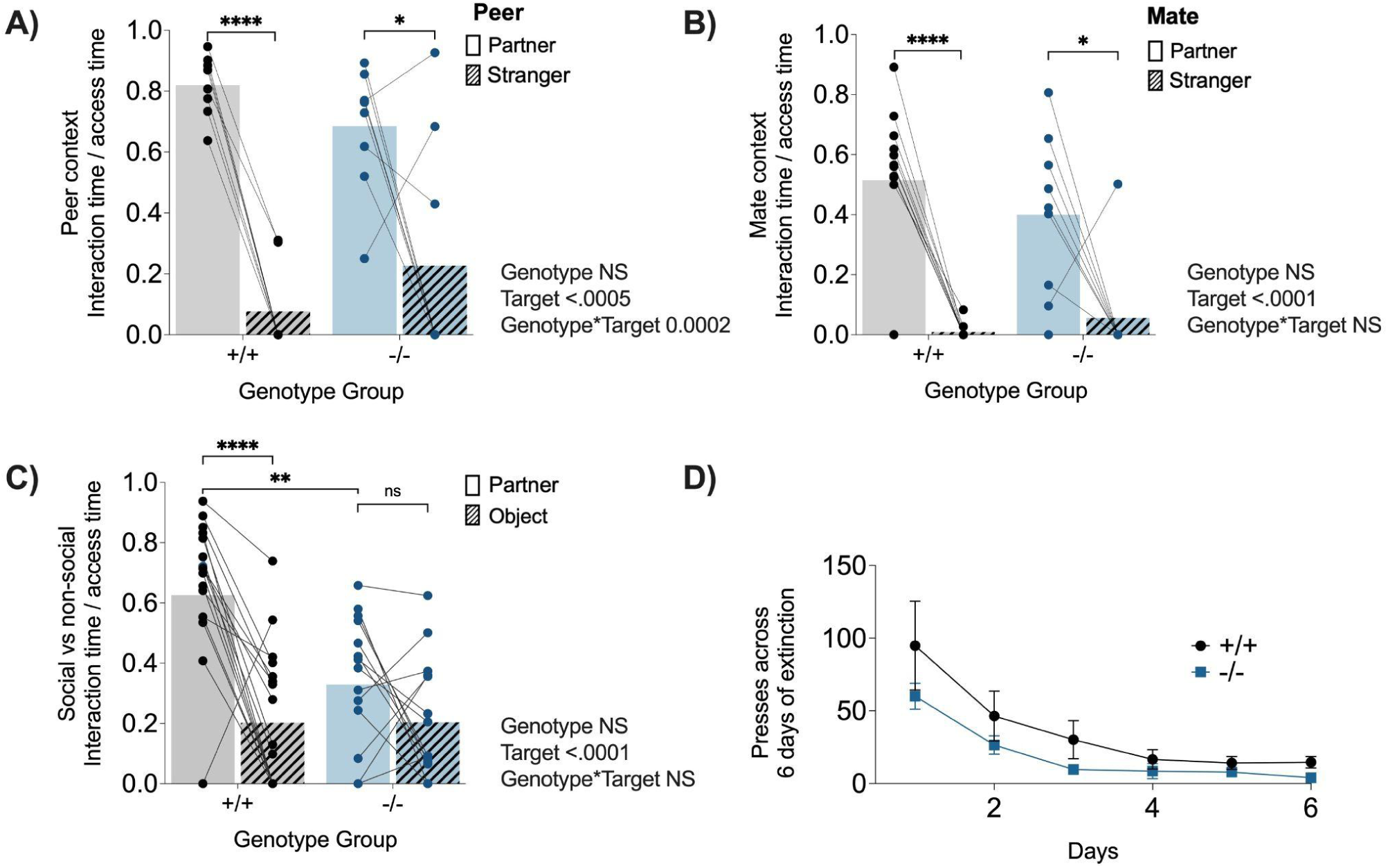
Reward interaction during operant testing (phases 1-3) and lever-pressing extinction curve. **A)** Even though *Oxtr*^1−/−^ animals did not show selective motivation for the peer partner versus the stranger reward context (Fig 3B), both groups spent more of their reward access time huddling with the partner than the stranger. Paired t-test: +/+ t(7) = 14.63, p = 0.000002; −/− t(8) = 2.85, p = 0.02148. **B)** In the mate partner vs stranger reward context (Fig 3C), both WT and *Oxtr*^1−/−^ voles spent more of their reward access time huddling with their mate partner than strangers. Paired t-test: +/+ t(11) = 6.649, p = 0.000036; −/− t(8) = 2.781, p = 0.02388. **C)** In the social versus non-social reward context (Fig 3D), WT animals preferred to spend a larger proportion of their reward access time interacting with the social stimulus than with the non-social stimulus compared to *Oxtr*^1−/−^ who showed no difference in interaction duration by reward type. Paired t test: +/+ t(17) = 5.493, p = 0.000040; −/− t(14) = 1.657, p = 0.1197. Overall, *Oxtr*^1−/−^ spent a lower proportion of reward access time interacting with their partner compared to WT. Student’s t test: t(11) = 4.3509, p = 0.0011.**D)** Average number of presses by genotype across 6 days of extinction testing with no lever actions. Repeated measures MANOVA revealed no difference in extinction between genotype groups F(1,7) = 0.3038, p = 0.1881.

**S4.**
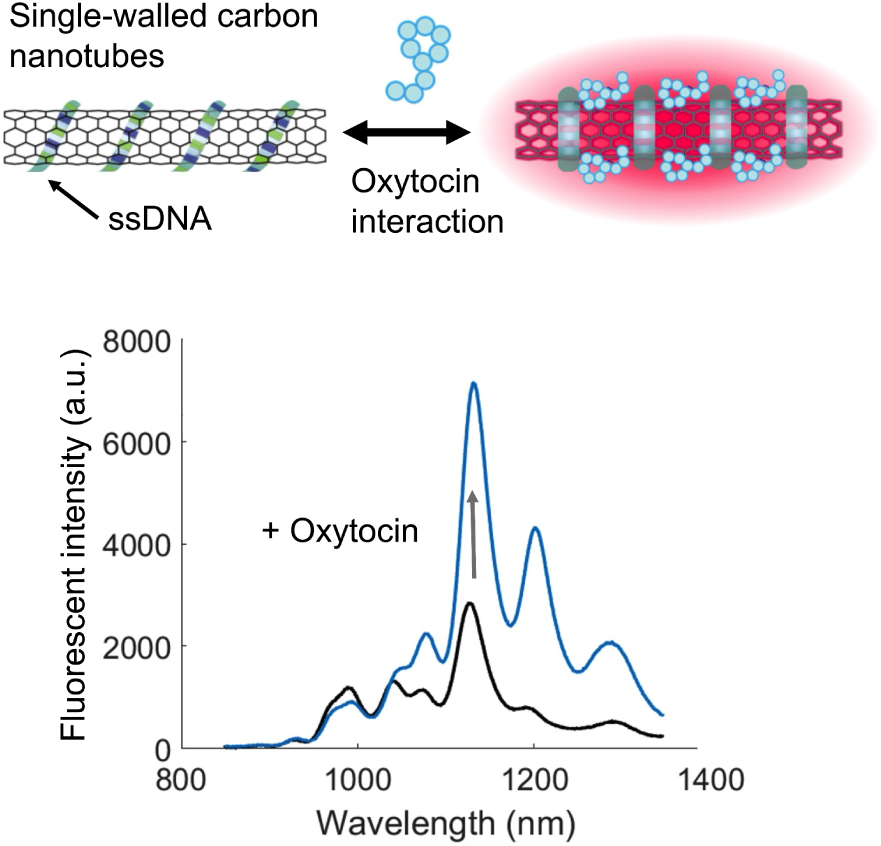
Mechanism of NanOx nanosensor. NanOx is based on single-walled carbon nanotubes functionalized by single-stranded DNA (ssDNA) that confers selectivity for oxytocin. A unique ssDNA sequence was identified to modulate the fluorescence intensity of single-walled carbon nanotubes in the presence of oxytocin.

**S5.**
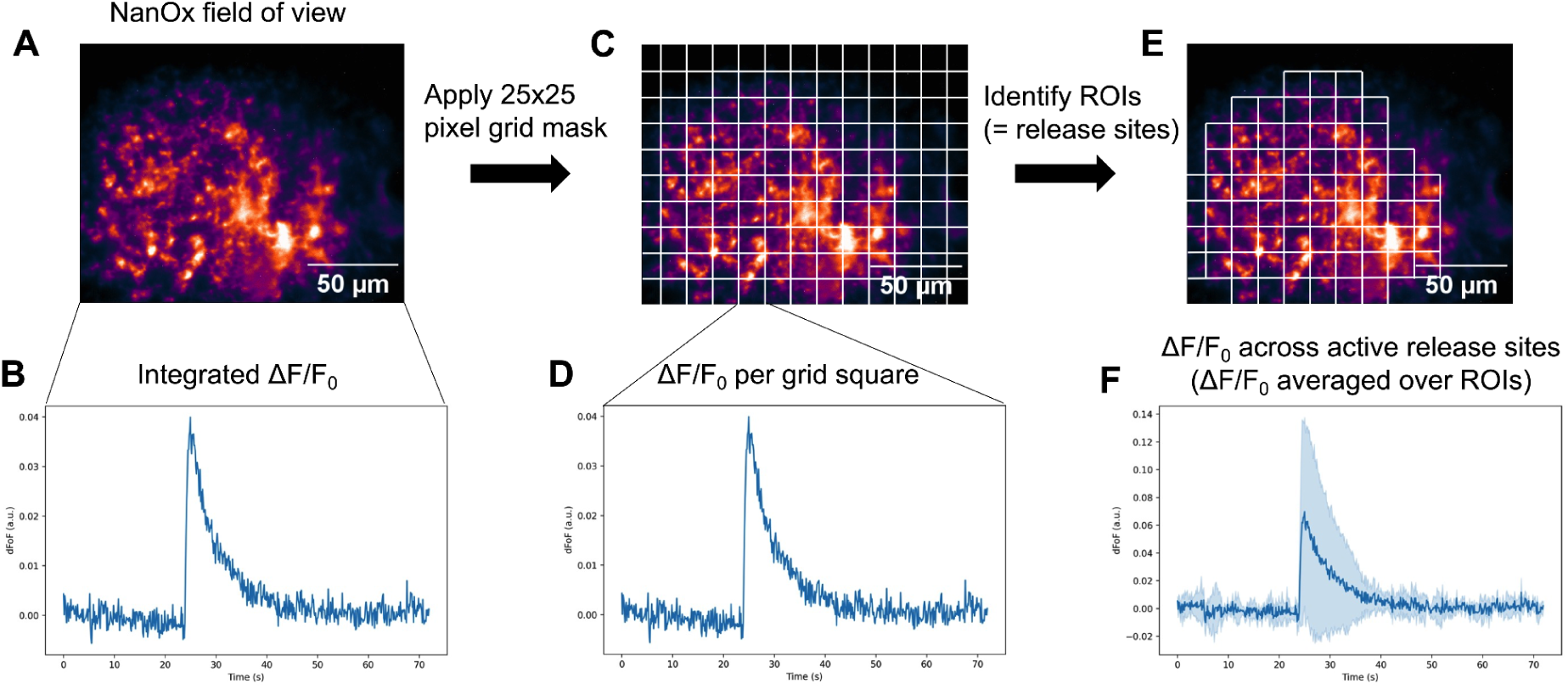
Image processing and data analysis of nanosensor fluorescence response. **A**) Imaging movie files to be processed showing the entire field of view (175 µm by 140 µm). **B**) Integrated ΔF/F_0_ (ΔF/F_0_ for the entire field of view) was calculated as ΔF/F_0_ = (F-F_0_) /F_0_, where F_0_ is the average intensity for the first 5% of frames and F is the dynamic fluorescence intensity. These data are subsequently averaged over three stimulation replicates. **C**) A 25×25 pixel (corresponding to 6.8 µm by 6.8 µm) grid mask was applied to the image stack. **D**) ΔF/F_0_ was calculated for each grid square. **E**) Grid squares were identified as regions of interest (ROI) if the F around time of stimulation (200 frames) is 3 standard deviations above the baseline F_0_ activity. This physiologically corresponds to active oxytocin release sites. **F**) ΔF/F_0_ was averaged over all identified ROIs, corresponding to ΔF/F_0_ across active release sites. The solid line represents mean and the shaded region represents standard deviations.

**S6.**
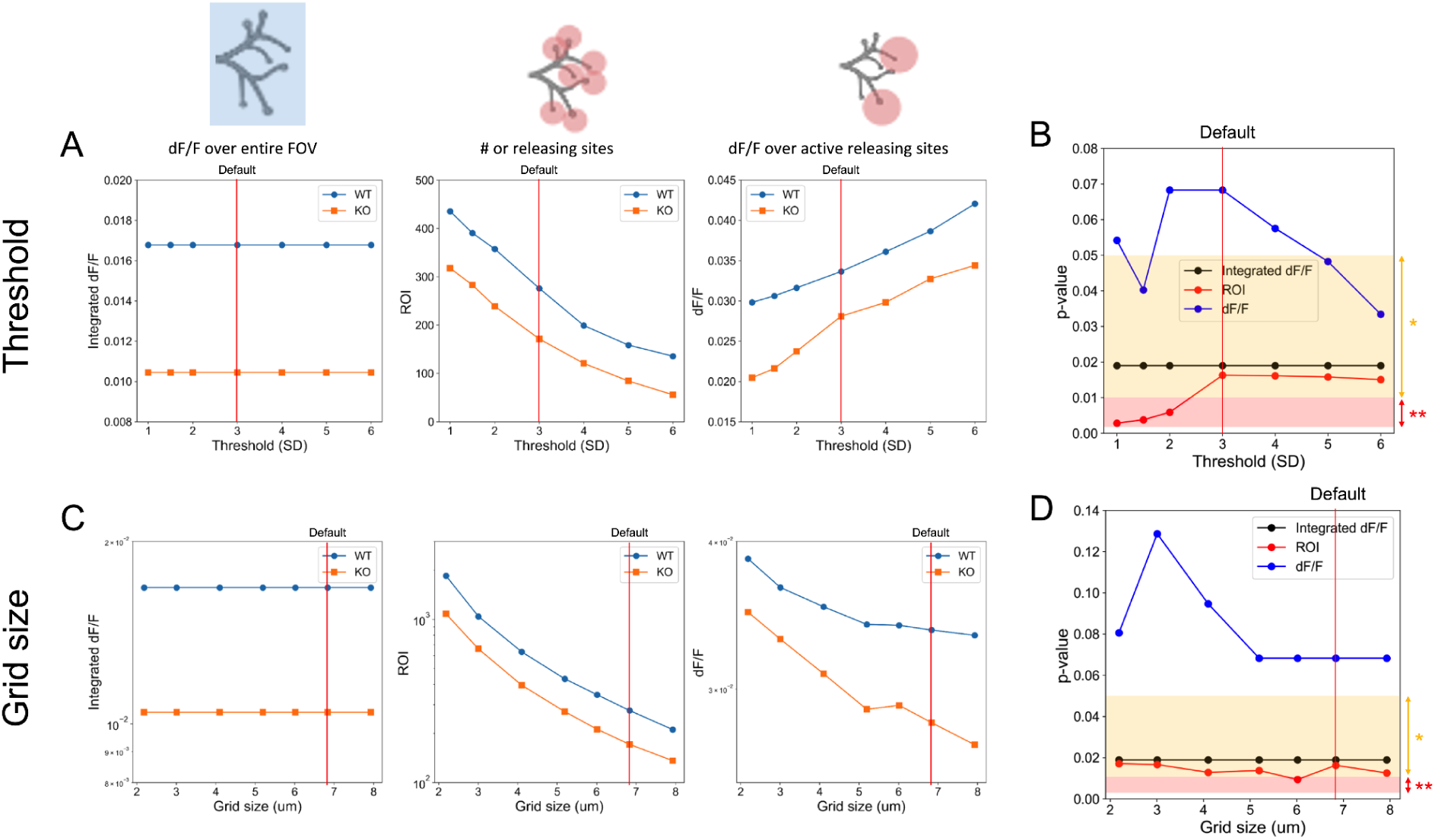
Threshold value and grid size dependence of the analyzed parameters. **A, B**) Grid squares are identified as regions of interest (ROI) if the F around time of stimulation (200 frames) is 3 standard deviations above the baseline F_0_ activity. We changed this threshold from 1 to 6 to examine how it changed each parameter (i.e., integrated ΔF/F_0_, ROI, and mean peak ΔF/F_0_) and the p-values comparing two genotype groups, while keeping the grid size as 25 × 25 pixels (corresponding to 6.8 × 6.8 µm). **A**) As expected, integrated ΔF/F_0_ does not change with changing threshold value (because the integrated ΔF/F_0_ analysis does not involve grid square analysis), more ROIs were identified with smaller threshold, and mean peak ΔF/F_0_ was lower for smaller threshold (because smaller threshold will include more spots with lower ΔF/F_0_). **B**) p-value for integrated ΔF/F_0_ (black), ROI (red), and mean peak ΔF/F_0_ (blue) as a function of the threshold value to identify active release sites. We picked 3 as the threshold value as p-value for ROI (red) was consistent from 3 onward. **C, D**) We changed the grid size from 8 × 8 pixels (corresponding to 2.2 × 2.2 µm) to 29 × 29 pixels (corresponding to 7.9 × 7.9 µm) to examine its influence on each parameter (i.e., integrated ΔF/F_0_, ROI, and mean peak ΔF/F_0_) and the p-values comparing two genotype groups, while keeping the threshold value as 3. **C**) As expected, the integrated ΔF/F_0_ does not change with changing threshold value (because the integrated ΔF/F_0_ analysis does not involve grid square analysis), more ROIs were identified with smaller grid size, and mean peak ΔF/F_0_ was higher for smaller grid size. **D**) p-value for integrated ΔF/F_0_ (black), ROI (red), and mean peak ΔF/F_0_ (blue) as a function of the grid size. We picked 25 × 25 pixels (corresponding to 6.8 × 6.8 µm) as the grid size as p-value for ROI (red) and for mean peak ΔF/F_0_ (blue) was relatively consistent after 19 × 19 pixels (corresponding to 5.2 × 5.2 µm), and this size is physiologically relevant considering the size of oxytocin neurons.

**S7.**
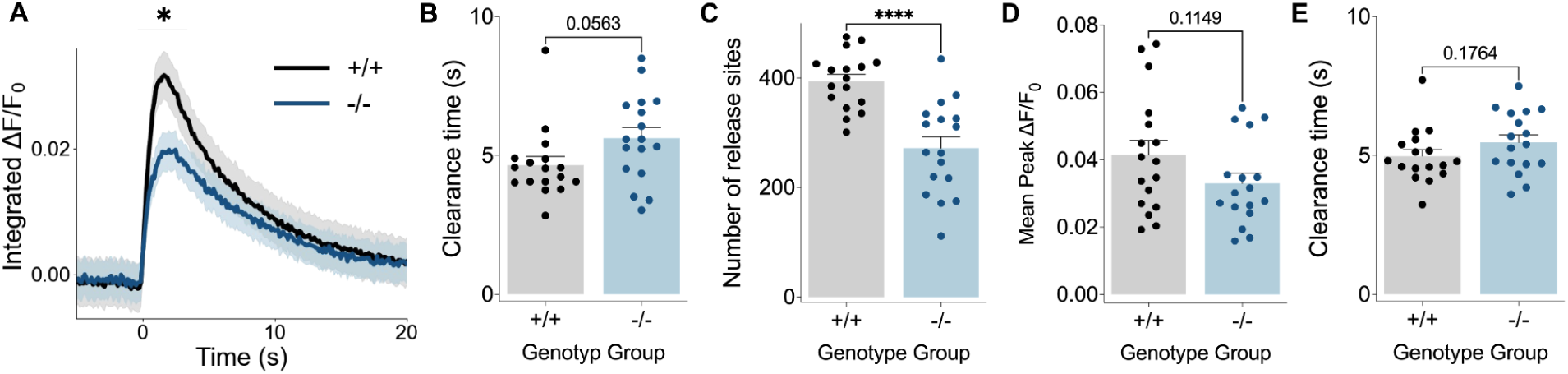
Fluorescence imaging with NanOx in nucleus accumbens before quinpirole application. **A**) Time-course of integrated ΔF/F_0_ in WT and *Oxtr*^1−/−^ voles. Student’s t-test: +/+ vs −/− – t(32) = 2.527, p =0.0166. **B**) oxytocin clearance time of integrated ΔF/F_0_. Student’s t-test: +/+ vs −/− – t(32) = 1.980, p =0.0563. **C**) The number of release sites identified in the field of view. Student’s t-test: +/+ vs −/− – t(32) = 5.085, p <0.0001. **D**) Mean peak ΔF/F_0_ values across identified release sites. Student’s t-test: +/+ vs −/− – t(32) = 1.621, p =0.1149. **E**) oxytocin clearance time averaged over identified release sites. Student’s t-test: +/+ vs −/− – t(32) = 1.383, p =0.1764. Data (mean ± SEM) from n = 10 voles and 17 slices for WT and n = 10 voles and 17 slices for *Oxtr*^1−/−^. * = <.05, ** = <.01, *** = <.001, ****<.0001 are determined by Student’s t test.

**S8.**
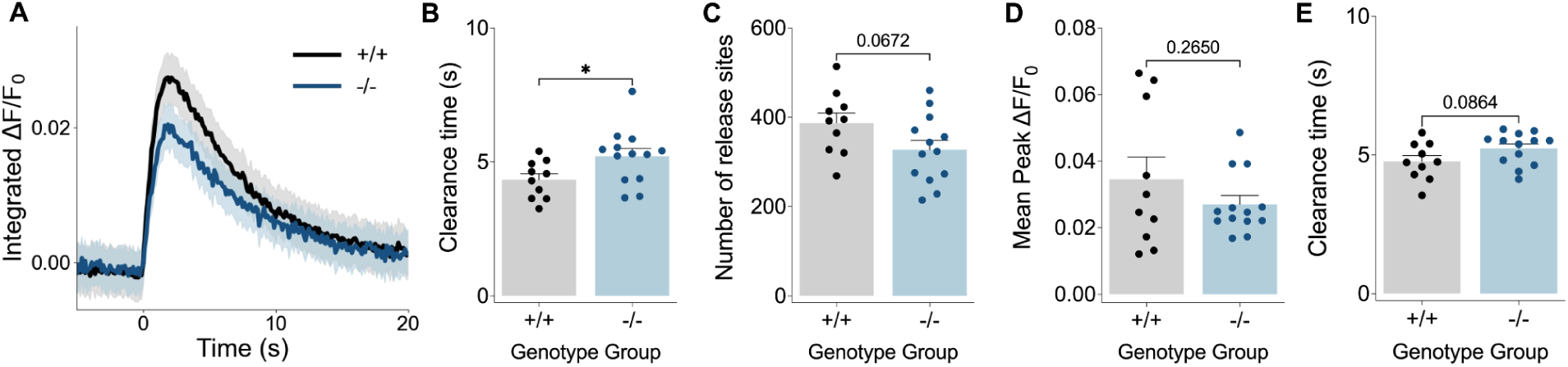
Fluorescence imaging by nIRCats reveals slower dopamine clearance and trends towards reduced release sites in the nucleus accumbens of OXTR mutant voles. **A**) Time-course of integrated ΔF/F_0_ in WT and *Oxtr*^1−/−^ voles. Student’s t-test: +/+ vs −/− – t(21) = 1.437, p =0.1654.**B**) oxytocin clearance time of integrated ΔF/F_0_. Student’s t-test: +/+ vs −/− – t(21) = 2.271, p =0.0338. **C**) The number of release sites identified in the field of view. Student’s t-test: +/+ vs −/− – t(21) = 1.930, p =0.0672. **D**) Mean peak ΔF/F_0_ values across identified release sites. Student’s t-test: +/+ vs −/− – t(21) = 1.145, p =0.2650. **E**) dopamine clearance time averaged over identified release sites. Student’s t-test: +/+ vs −/− – t(21) = 1.799, p =0.0864. Data (mean ± SEM) from n = 6 voles and 10 slices for WT and n = 6 voles and 13 slices for *Oxtr*^1−/−^. * = <.05, ** = <.01, *** = <.001, ****<.0001 are determined by Student’s t test.

