## Supplementary figures and images for "Oxytocin receptor absence reduces selectivity in peer relationships and alters neurochemical release dynamics in prairie voles"

### S1

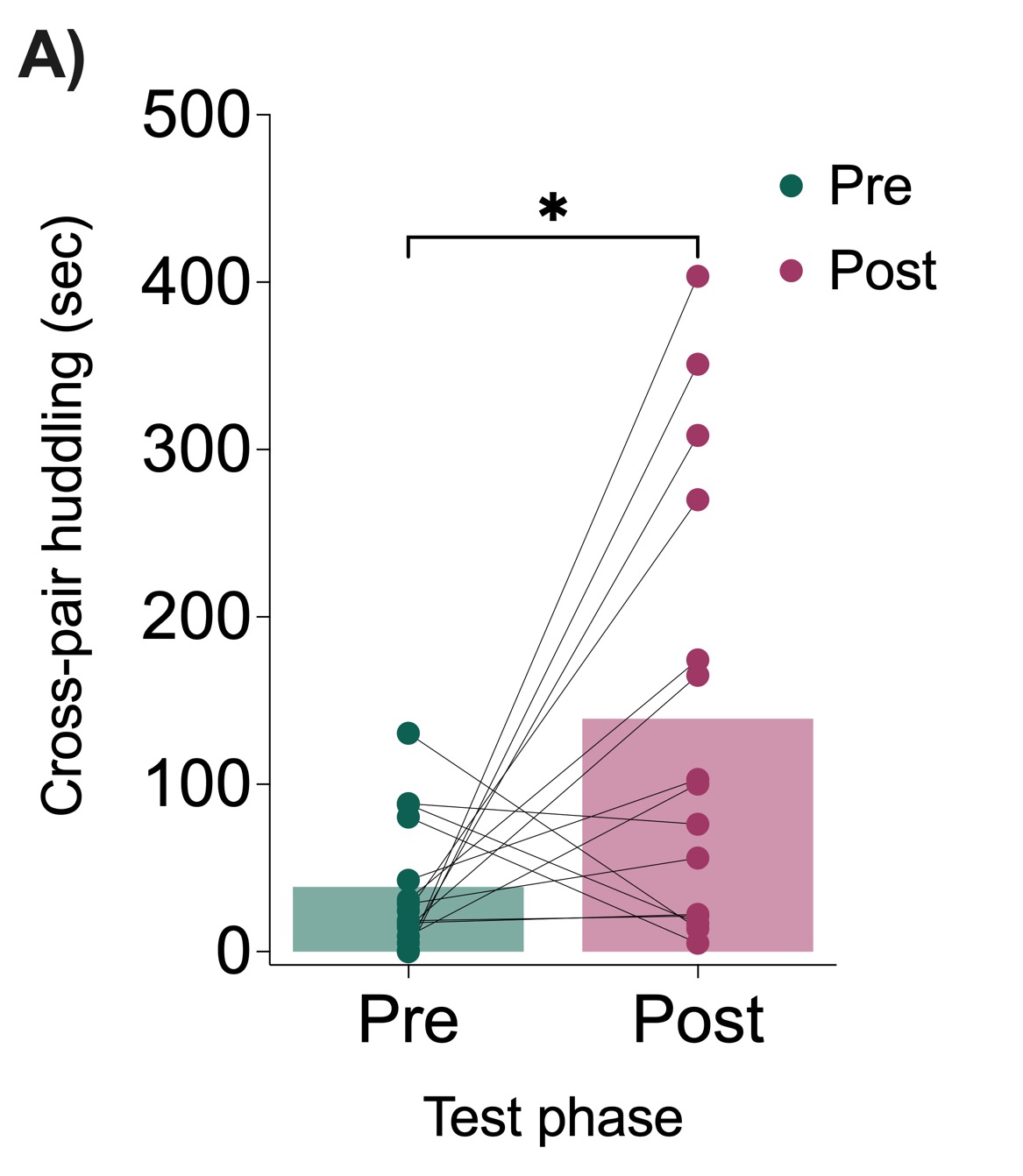

### S2

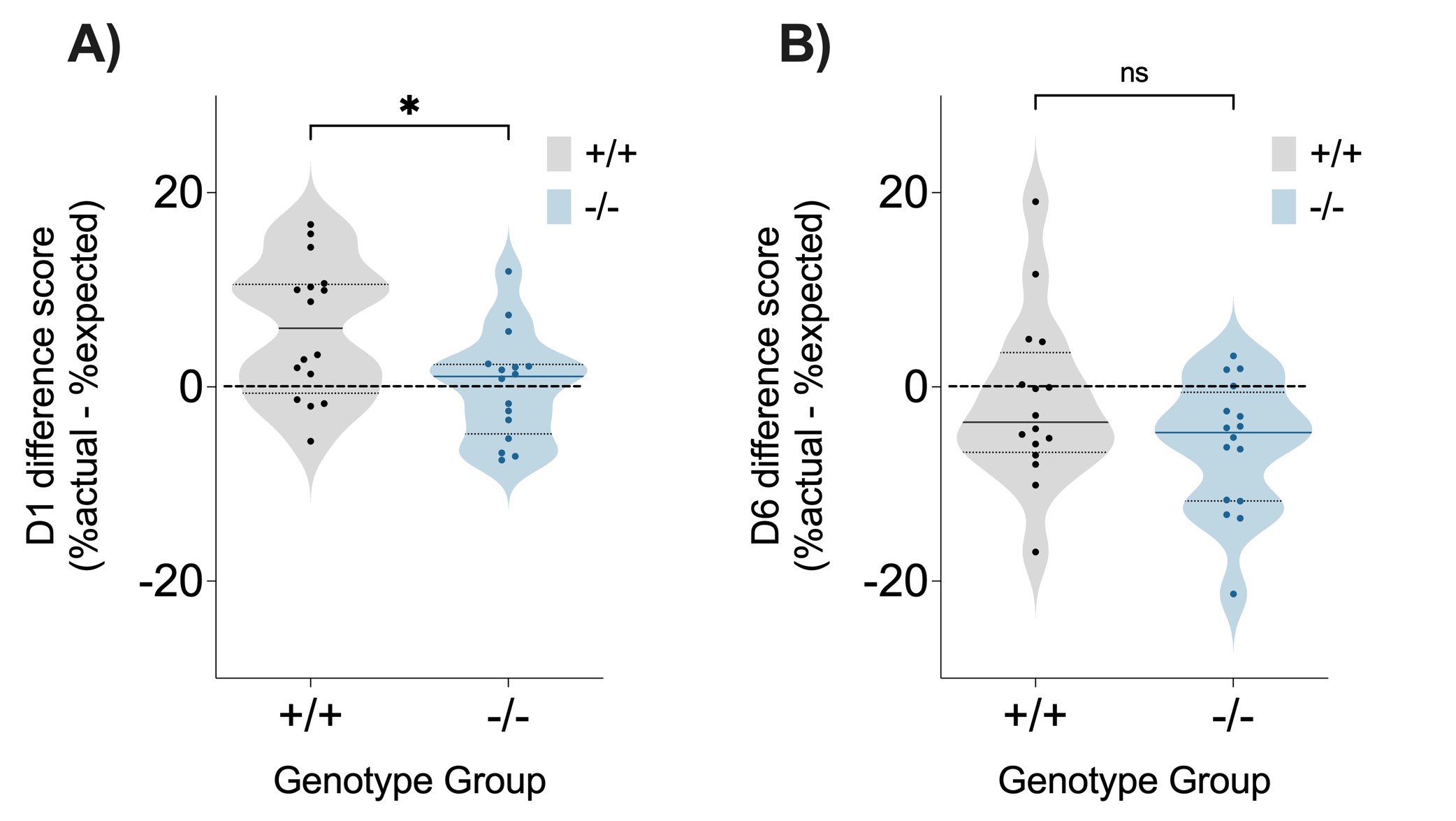

### S3

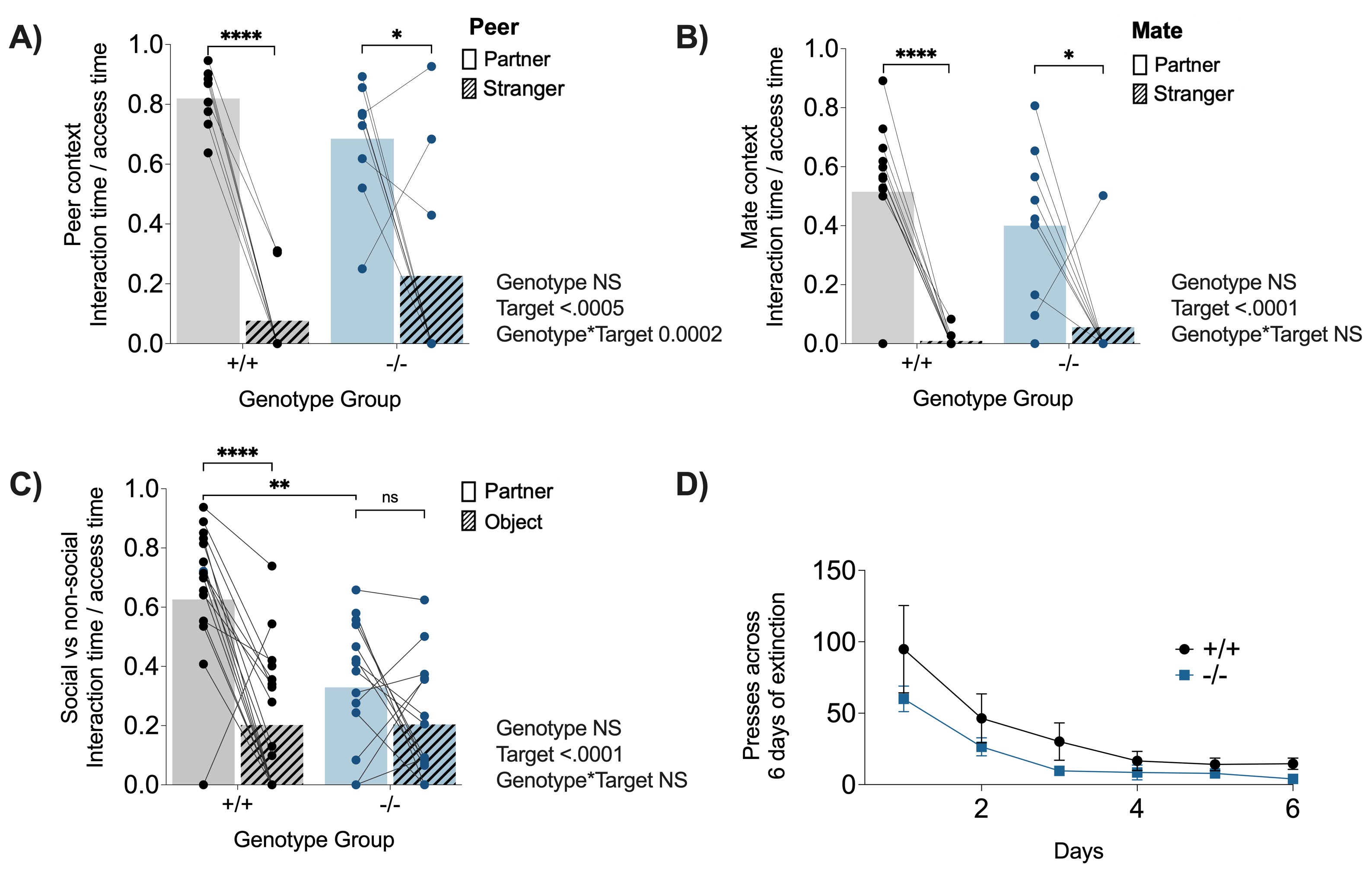

### S4

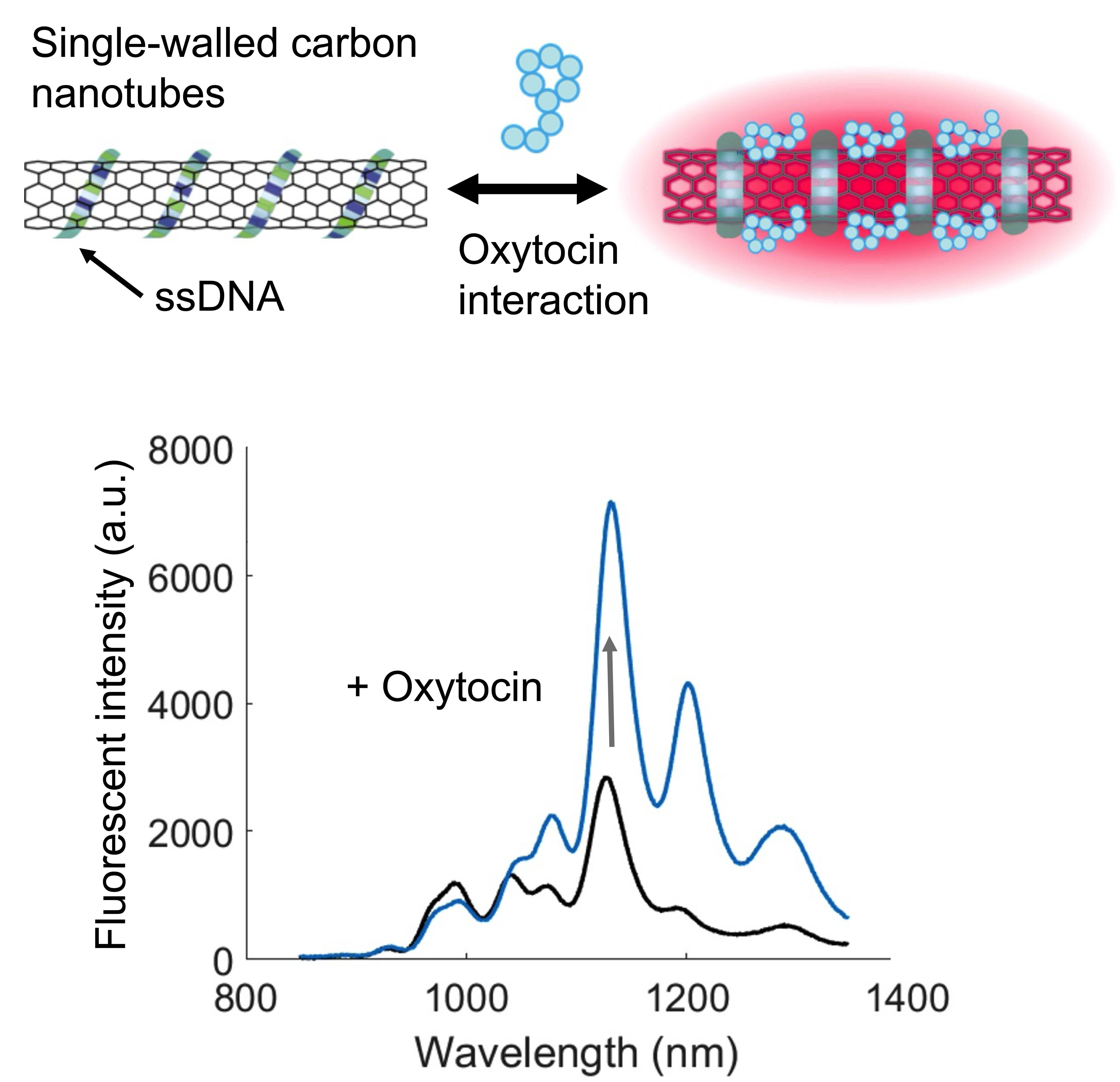

### S5

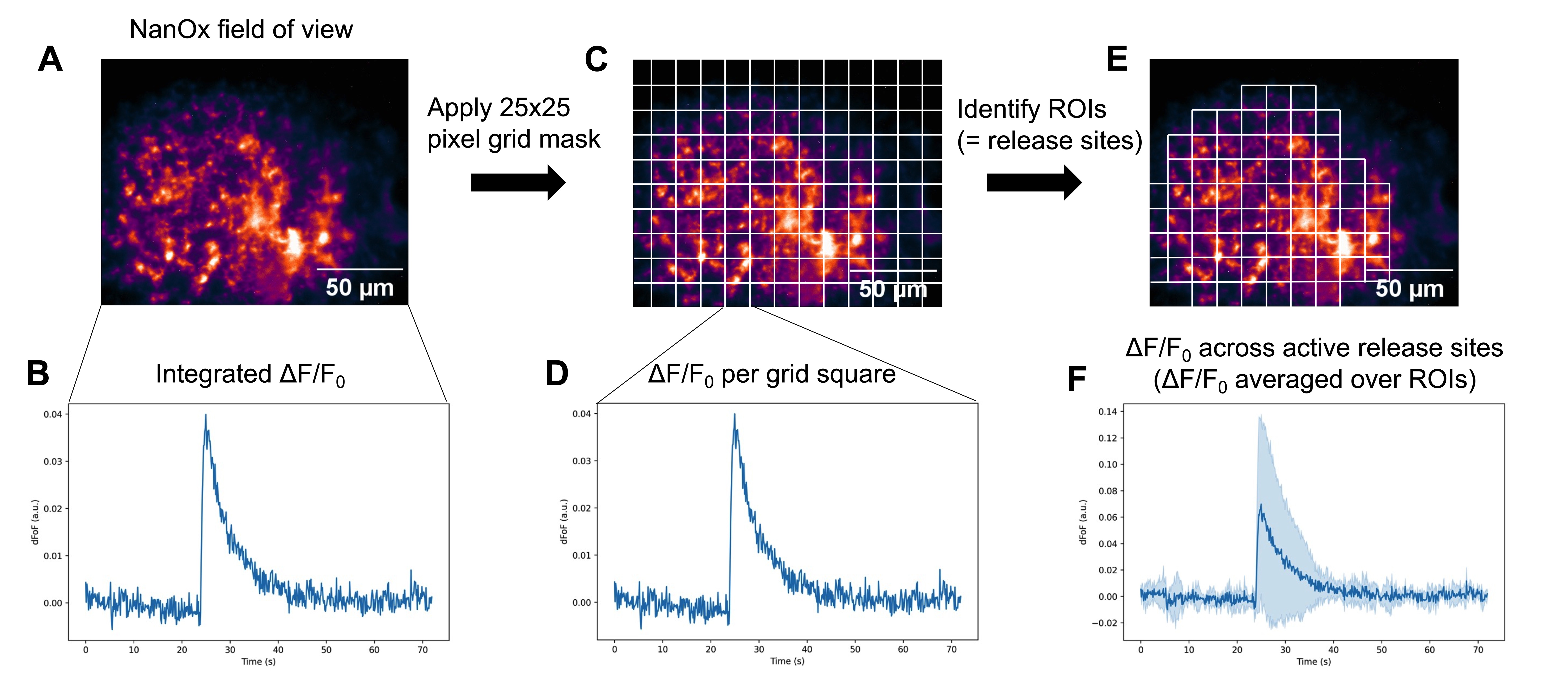

### S6

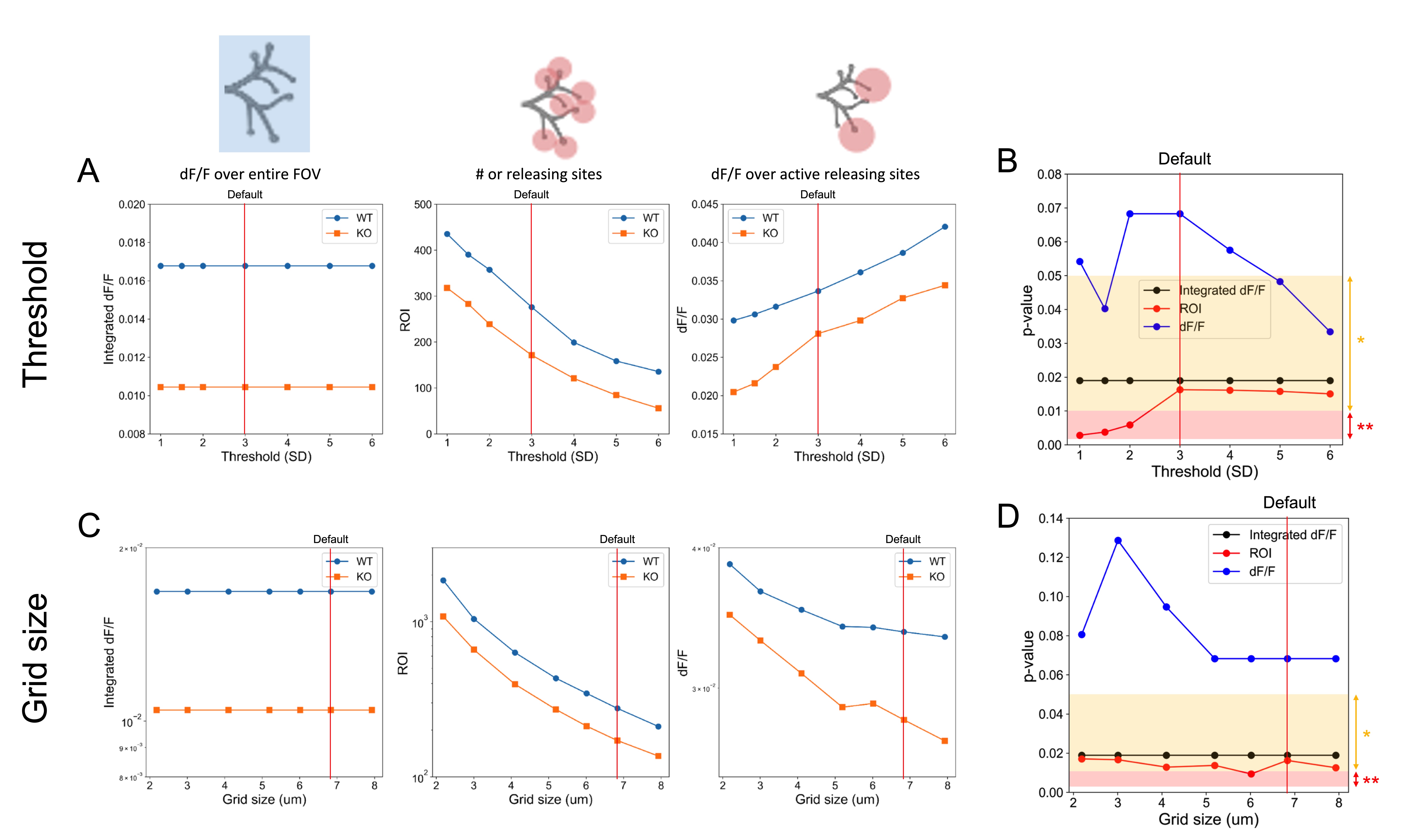

### S7

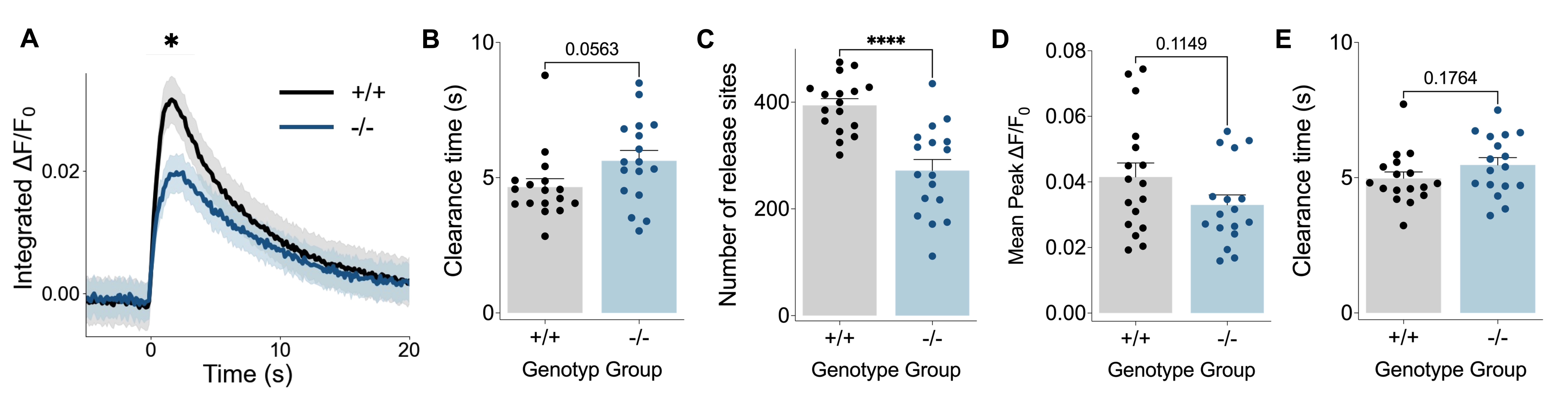

### S8

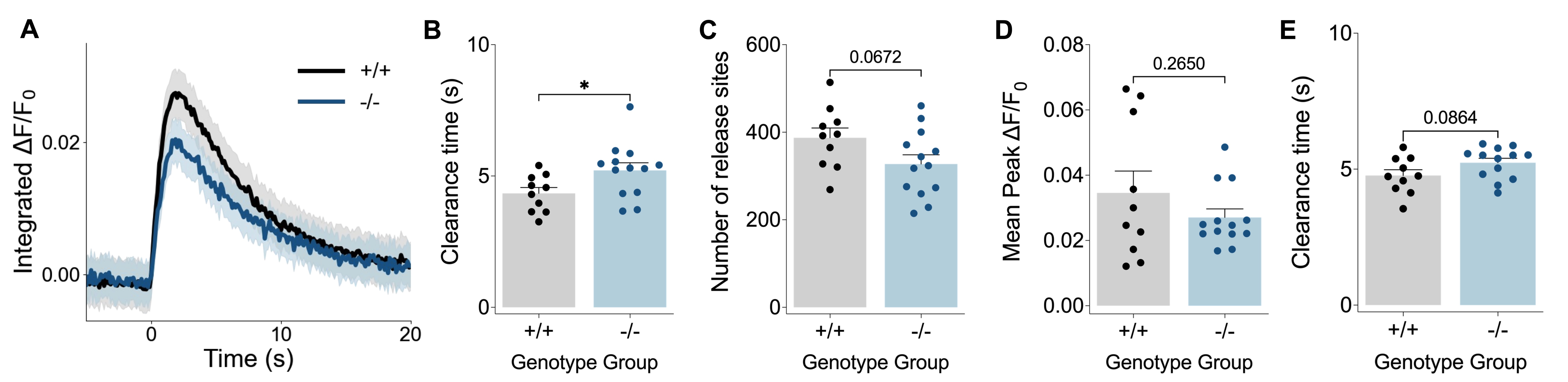
